# Transforming macromolecular structures into simulations of self-assembly with ioNERDSS

**DOI:** 10.64898/2026.01.27.702082

**Authors:** Yue Moon Ying, Mankun Sang, Gabriel Au, Smriti Chhibber, Yufeng Du, Jonathan A Fischer, Samuel L Foley, Sikao Guo, Ian Herzog-Pohl, Zixiu Liu, Hannah Roscom, Hassan Sohail, Sho S Takeshita, Margaret E Johnson

## Abstract

Macromolecular self-assembly is a fundamental process in living and engineered systems, producing molecular machines like the ribosome or highly symmetric viral capsids. Thanks to sources like the Protein Data Bank (PDB) and AlphaFold3, the final target complexes are often known, but these static structures do not provide information on the self-assembly process directly. Computational models provide critical tools to study these essential pathways of self-assembly, but substantial coarse-graining of assembly subunits is necessary to achieve computational tractability of these relatively slow processes while retaining multi-valency. While rule-based or local interactions overcome the often-prohibitive enumeration of all possible assembly intermediates, they must ensure global structural constraints are met. We here demonstrate ioNERDSS, a user-friendly Python package that transforms 3D atomic structures into coarse-grained models for immediate simulation with the stochastic reaction-diffusion NERDSS software, converting static structures into time-resolved assembly trajectories. NERDSS uses rule-based interactions to simulate multi-component self-assembly at the minutes timescales and without limits to complex size or growth pathways. With ioNERDSS, each protein chain is defined by a rigid subunit with discrete interfaces and explicit orientational constraints that enforce a structured assembly. Repeated subunits (such as in viral capsids) are regularized to preserve the target topology across distinct stochastic assembly pathways, supporting assembly of structures with thousands of subunits. We initialize pairwise binding affinities using open-source machine-learned prediction tools, and our default coarse-grained (CG) models are all constrained by thermodynamic reversibility to reach an equilibrium steady-state. The binding rates and subunit abundances necessary to perform simulations are initialized at default values but represent the key variables (along with affinities) that cells and thus users would tune to control productive assembly. Benchmarking on over 40,000 PDB structures shows that the majority of CG models stochastically assemble into target structures. The ioNERDSS Python library links directly to open-source tools for visualization and analysis to facilitate fast and user-friendly structure validation and analysis of output for thermodynamic, kinetic, and nonequilibrium drivers of macromolecular self-assembly.

## 1. Introduction

From vesicle trafficking and protein translation to DNA transcription, living systems rely on multi-subunit macromolecular complexes to function(1, 2). The assembly of the ribosome (>60 subunits, all unique) or the HIV-1 Gag lattice (>2000 subunits)(3, 4) requires many successful pairwise association events of monomers into units with correct stoichiometry in the nonequilibrium environment of the cell. While the biogenesis and assembly of ribosomes have been studied in detail (5, 6), the assembly pathways used by the hundreds of other macromolecular complexes(2) often lack systematic characterization. One factor limiting quantitative measurements of assembly pathways, kinetics, and control mechanisms is access to efficient and structurally accurate models that provide time-resolved simulations to help interpret experiments. The structures of the final assembled complexes are typically no longer unknown, thanks to the protein data bank (PDB) (1) and the predictive powers of AlphaFold3 (7). Coarse-grained (CG) models offer a computationally tractable and structurally faithful approach to study mechanisms of macromolecular assembly, but it is often a laborious process to construct a correctly assembling CG model retaining geometric features and reasonable thermodynamics from known 3D molecular structures. Currently, no tools convert 3D macromolecular structures directly into tractable CG models for multi-valent self-assembly trajectories. To address this, we present ioNERDSS — a “structure-in-simulation-out” Python library that automates coarse-graining from atomic or PDB-level structural data to rate-based CG models. In doing so, ioNERDSS bridges a gap between detailed structural information and mesoscale assembly dynamics, making CG self-assembly tools accessible to scientists in structural and cell biology, with applications in synthetic biology and materials design. The parameterized models and CG subunits are immediately executable with the NERDSS software for stochastic, rigid-body reaction-diffusion (RD) simulations (8), and for small assemblies a corresponding set of deterministic rate equations. The automated ioNERDSS pipeline supports exploratory modeling, classroom demonstration, and rigorous hypothesis testing without requiring extensive computational expertise.

A key benefit of using physics-based computer simulations is that from a target assembled structure, the number of possible pathways to reach this structure is vast and entirely underdetermined from just the structure alone. ioNERDSS generates an initial model that obeys reversible interactions between all contacts to eventually produce an equilibrium steady-state. This steady-state theoretically includes the target structure, but due to thermodynamic and/or kinetic effects, it may be improbable or impossible to reach this state from initial monomeric subunits on a reasonable (minutes-hours) timescale (9). However, a significant advantage of the rate-based models that are defined and simulated by ioNERDSS, including NERDSS (8), in contrast to energy-function based methods like Molecular Dynamics (MD), is that they are relatively efficient and scalable (10-13), allowing this vast design space to be feasibly explored. Pathways of single-component assemblies like filaments (14) or viral capsids (15-17) are fairly well-studied using these models, but systematic applications to structures built from a variety of distinct subunits, similar to many complexes in the PDB, is relatively recent (9, 18-28). These studies have collectively demonstrated how the *process* of self-assembly can dramatically impact the practical yield of functional complexes and target structures compared to what one would naively expect from purely thermodynamic considerations. For example, a fundamental barrier to productive self-assembly stems from partially assembled kinetic traps, with one of the simplest forms being monomer starvation. Intermediates along the proper assembly pathway form (e.g. dimers) but cannot directly transition to the completed complex (two dimers cannot form a trimer), and waiting for disassembly can create an exponentially growing time-delay with increasing assembly size(9). Rate-based models have established that the topology and number of subunits of macromolecular complexes are key factors in determining how hard self-assembly can be(9, 29). However, beyond these purely structural factors, a vast design space exists to achieve controlled and productive self-assembly for any structure by creating hierarchies in binding rates (9), subunit stoichiometries(22), interaction energies(27), or subcomplex rates(9, 29).

Exploring assembly pathways and equilibria using rate-based models must overcome an inherent challenge: the possible states of subunits must be pre-specified, with full enumeration(9, 16, 30) or rule-based approaches(31-34) thus an essential step in model set-up. Full enumeration of all subcomplexes or intermediates scale exponentially with assembly size and therefore becomes prohibitive for large assemblies. Rule-based methods circumvent enumeration by requiring only pairwise interactions, enabling models with highly diverse substates(32-34). However, without a spatial representation of multi-valent subunit structure, rules on their own cannot encode orientational constraints that lead to self-limiting assemblies and well-defined topologies (e.g. trimer, proteasome, or capsids(16)) or capture steric exclusion that prevents unphysical overlap(35). Although coarse molecular structure in other rule-based RD methods support assembly of disordered structures(36) and polymers(37), here we focus on applications to the finite-sized macromolecular complexes that dominate the PDB, and self-limited assemblies like cages and filaments(4, 9, 13, 38, 39). The NERDSS software(8) combines rule-based interactions with enforced orientational constraints to ensure these structured topologies are realized without limits to assembly size or pathways, and ioNERDSS computes these constraints directly from PDB/CIF files to automate model set-up. For small enough assemblies (<12 subunits), it also automatically enumerates assembly pathways to generate integrable non-spatial rate equations that include only geometrically admissible intermediates, reproducing the steric constraints of the spatial model.

Beyond subunit structures and orientations, therefore, a key step in model set-up by ioNERDSS is the specification of binding association and dissociation rates, *k*_on_ and *k*_off_. Inferring the values of these rate constants from experimental time-resolved data is a primary application of the ioNERDSS pipeline; most of these rates are not known experimentally (although databases exist(40)) and directly predicting rates purely from 3D structures is not currently possible with deep-learning-based methods. To initialize a fully executable model, we therefore simply assign association rates closer to the diffusion-limit (7×10^7^ M^-1^s^-1^) and use deep-learning methods(41) to instead predict the binding affinities *K*_D_ (or equivalently binding free energies Δ*G*) for each pairwise contact in the structure. Given *K*_D_ and a defined *k*_on_, we then uniquely define the dissociation rates using the well-known relation *k*_off_ = *k*_on_*K*_D_. In this way, ioNERDSS does provide structure and sequence-informed constraints(41) on the predicted thermodynamic equilibrium of the assembly, albeit subject to the simplifying assumption that a pair of binding interfaces has the same Δ*G* whether between monomeric subunits or between assembled intermediates. Importantly, this assumption can be relaxed in NERDSS by specifying conditional, state-dependent rates for any interface pair to support active and allostery-induced changes to affinities or rates during assembly (8).

Although models built with ioNERDSS are intended for rate-based simulations rather than MD, the software adopts the same user-friendly workflow common in the thriving ecosystem of MD simulation for building systems and setting up force fields directly from PDB structures(42, 43). Because all-atom MD is far too expensive to simulate multi-component self-assembly from monomeric building blocks, the only MD methods capable of studying assembly pathways comparable to rate-based methods are ultra-CG MD (44) (45-48). The advantage of the ultra-CG MD approach for studying self-assembly is having the Δ*G* between any components emerge naturally from sequence and structure of proteins (44), with built-in mechanics supporting polymorphism (45-48) and flexibility (45-48). While all-atom models offer automated setups, deriving components and force-fields for ultra-CG MD from 3D atomic structures typically requires substantial user expertise(44, 49-52). In contrast, ioNERDSS provides a more user-friendly model setup. Ultra CG MD is also still significantly less efficient than RD or other rate-based approaches, which support straightforward tunability of pairwise binding kinetics to map directly to emergent experimental timescales (38, 53, 54). Ultimately MD simulations offer a ‘bottom-up’ approach to parameterize the more efficient rate-based RD models through simulations of pairwise association. While this would offer a powerful route to parameterize our self-assembly models, currently the enhanced sampling required to extract either binding rates (55, 56) or binding free energies (57-61) for each protein binding pair still requires heavy compute and user expertise (62, 63), rendering it infeasible for the ioNERDSS pipeline. Accessibility and ease-of-use is an important feature of ioNERDSS to enable efficient and quantitative self-assembly of multi-component systems by nonexperts.

In this paper, we describe the steps necessary to automate the coarse-graining of macromolecular complexes for self-assembly, with several additional steps required for repeated subunits to ensure correct topology of assembled structures. By defining coarse-grained RD models and ODE systems for the same structural specification (for smaller complexes), ioNERDSS enables users to compare spatial, stochastic behavior with well-mixed deterministic dynamics under identical thermodynamic and kinetic parameters. We demonstrate several example applications of ioNERDSS and we detail when ioNERDSS can fail to generate correct assembly models when repeated subunits exhibit high variation between contacts and inter-molecular orientations, based on systematic benchmarking of >44,000 PDB multi-protein structures, with Python protocols provided for structure validation and troubleshooting. We provide a platonic solid library for simple design of ideal structures with easily tunable sizes. To streamline usage, ioNERDSS can be directly imported as a Python library, allowing users to call its functions without interacting with the source code, including analysis of simulation output and outputs to open-source visualization tools including Simularium (64) and OVITO (65) for interactive, shareable 3D visualization of simulation trajectories. Finally, we provide an online server for easy structure-to-simulation trials at nerdssdemo.org.

## 2. Methods

The ioNERDSS library provides a comprehensive pipeline for processing Protein Data Bank (PDB) structures and converting them into coarse-grained molecular models ready for running NERDSS simulations. Other functionalities include ODE-based kinetic models, platonic solid templates for symmetric assemblies, and a library of analysis tools for NERDSS output.

### 2.1. Converting PDB structure to ready-to-run NERDSS model

We use the Biopython package(66) to read the PDB or mmCIF files and identify *N*_chain_ protein polypeptide chains, ignoring all non-protein components and non-standard amino acids. The ioNERDSS pipeline identifies *N*_mol_ distinct proteins in the assembly, where *N*_mol_ *N*_chain_ . For example, for the HIV Gag lattice (PDB 5L93), the assembly file contain*s N*_chain_ = 18 but produces *N*_mol_ =1 distinct protein type. For each of the chain*s i* ∈ 1: *N*_chain_, we first compute (i) a geometric center of mass, 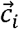 (COM; unweighted) and (ii) a molecular radius *R*_*i*_ (root-mean-square distance of atoms from the COM) which is needed to assign translational diffusion coefficient *D*_t_ and rotational diffusion coefficient *D*_r_ (assuming isotropy) to each subunit using Einstein-Stokes relation (SI 1.3). To determine interactions between molecules, candidate chain–chain contacts are prefiltered by axis-aligned bounding boxes to avoid redundant iteration over distant chain pairs. Residue-level contacts are then found by *k*-dimensional tree (KD tree) queries over Cα coordinates within a distance cutoff *r*_cut_ = 6 Å . A chain-chain contact is accepted as a *bona fide* interaction only if *each* chain contributes at least *N*_cut_ =3 contacting residues, to eliminate incidental contact between chains that is unlikely to stabilize an unbound to bound transition. These default hyperparameters can be user-modified (Table S1). We assume that there is at most one interaction between any pair of chains, and therefore each interface in the interaction is coarse-grained to an interface point placed at the geometric center of all the contacting Cα coordinates, denoted by a vector 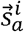 for subunit *i* and interface *a*. Because of the one interaction per pair limit, a chain pair (*i, j*) maps uniquely to an interaction pair (*a, b*) The norm of the vector between the two interacting sites, 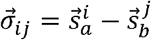, gives binding radius 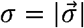. The orientations between the CG protein subunits *i* and *j* are first constrained by the vectors 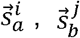, and 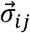 to define two rigid body angles *θ*_1_, *θ*_2_. An additional three dihedral angles, *ϕ*_1_, *ϕ*_2_ and *ω*, are required to uniquely constrain the rigid-body orientation of one CG subunit relative to the other (with some exceptions for perfectly colinear vectors), requiring body-fixed reference coordinates that are assigned to each subunit per interaction 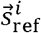. Between every two protein subunits *i* and *j* in the complex, there are therefore either zero or one pair of interfaces, with a total of *N*_edge_ interactions in the assembled structure (e.g. each chain is a node, and each interaction is an edge). If there are no repeated subunits in the assembly (*N*_mol_ *= N*_chain_), no further modifications are made to subunit CG structures or each set of interaction orientational constraints per *ij* pair 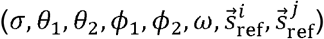, producing *N*_rxns_ = *N*_edge_ reversible reactions.

### 2.2 Repeated chains require extra refinement in ioNERDSS

If the macromolecular complexes contain *N*_rep_ copies of a protein chain, then ioNERDSS must perform additional steps to regularize the structure and interactions of these cases. This is because each subunit must be mapped to a single rigid-body geometry and a single set of pairwise reactions, regardless of how many copies are in the assembled structure. Repeated chains within complexes typically have the same contacting partners as one another with small thermal variations in the positions of interfaces, and thus we can simply pick a structure from *N*_rep_ copies and assign this structure uniformly to all the copies. However, we note that if a repeated chain interacts in a distinct molecular ‘neighborhood’, this requires more care (see results below). We identify repeated chains using polymer-entity IDs from the mmCIF header. If the file is in raw PDB format or these IDs are not present, we use sequence alignment in the Biopython package, with a similarity score cutoff of 0.5. It is important that we identify distinct vs repeated subunits, as distinct subunits have independently tunable copy numbers and affinities that can strongly influence assembly pathways. Structural homology is therefore not relevant. A single unique set of interfaces for a repeated subunit is defined primarily based off the CG positions of the interfaces and their partner sites, but residues in the interfaces can also be used to test for steric clashes in special cases (see SI). There are three general types of interface interactions. The heterodimeric case is the simplest (A+B), where both protein subunits are distinct, and therefore the interfaces are also unique to their subunits, 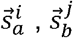. There are two types of homodimeric interface contacts, heterotypic and homotypic. For heterotypic interactions, such as in the actin polymer which binds head-to-tail, the subunits are the same, but the interfaces are different, 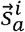 binding to 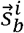. For homotypic interactions, the subunits and the interfaces are the same, or 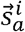 binding to 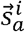. We annotate whether these homodimer interactions are homotypic or heterotypic during these steps, as they can produce distinct stoichiometries and reaction parameters. In particular, we note that in some special cases of usually idealized subunits like in platonic solids or user-defined subunits like the clathrin trimer (which is actually 6 polypeptide chains), these subunits can contain repeated copies of the same interface. Despite carrying the same binding properties, each of these interfaces must have a distinct label (e.g. *a*.*b*.*c*) to go along with their distinct coordinates. It would be erroneous to assert that each leg in a clathrin trimer has only a single leg on another trimer it can interact with (e.g. *a* only binds *a*), hence the need to identify structural and residue-level signatures of repeated interfaces and interactions (see SI Methods). Having established a unique set of interfaces for this subunit, the template subunit geometry for *i* = 1 is then aligned using rigid-body Kabsch transforms to the remaining *N*_rep_ − 1 copies to replace their existing coordinates with an identical rigid-body subunit, generating a newly regularized assembly.

From this newly regularized assembly we re-evaluate each set of interaction orientational constraints per *ij* pair 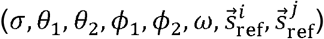, producing N_rxns_ <N_edge_ reversible reactions. With repeated interfaces (e.g. clathrin trimer), we enumerate and write out all possible reaction pairs using that interface due to the distinct labels per interface. Even after the previous subunit regularization step, there can be cases where the sets of interaction orientational constraints per repeated interactions produce discrepancies, particularly in angles. We then calculate a simple average for bond length and bond angles (*σ, θ*_1_, *θ*_2_) and a circular mean with special handling for high deviation for dihedral angles (*ϕ*_1_, *ϕ*_2_, *ω*). ioNERDSS writes out one molecule file (.mol) per CG molecule type and a parameter file (.inp) with molecule copy numbers, simulation parameters, and all reactions with their parameters, which are ready to run with the NERDSS software. If repeated subunits are arranged in a regular sphere (like the virus), we can exploit that radial symmetry to ensure a single curvature is produced during stochastic assembly by positioning all rigid subunits at a single radius from the sphere center(4, 67) (see hyperparameter options in SI Table S1).

### 2.3 Assigning reaction rates for each interaction pair

By default, we simply set the binding free energies per pairwise interaction to Δ*G*_ref_ = -16*k*_B_*T*, where Δ*G* = *G*_bnd_ − *G*_unb_. However, a more powerful approach that we use on the online webserver and illustrate in tutorials is to import and call the ProAffinity-GNN Python software to predict binding free energies, Δ*G*_GNN_ (41). ProAffinity-GNN estimates interface binding affinities by combining structure aware graph attention neural networks with protein language model embeddings, trained specifically for protein-protein interactions (41, 68). Here, ioNERDSS integrates a simple and user-friendly implementation that does not require canonical FASTA sequences for each binding partner to align/renumber PDB chains, because it is faster and more portable to evaluate. For our calculations, sequences are reconstructed directly from the observed residues in the PDB files (converted to PDBQT using AutoDockFR (68)), and ESM-2 embeddings (69) are computed on these structure-derived sequences. This inference-only path prioritizes ease of use at the cost of a modest reduction in accuracy relative to the full pipeline with FASTA sequences (see SI 3 for details). The binding free energy only constrains the ratio of rates, *K*_D_ = *c*_0_exp (Δ*G*/*k*_B_*T*)= *k*_b_/*k*_a_, with *k*_B_ Boltzmann constant, *T* the temperature, and the standard state *c*_0_ = 1M. With no reliable tool (yet) to rapidly predict association and dissociate rates, the association rates for all reactions are assumed to be closer to but below the diffusion-limit (*k*_a_ = 7.2 ×10^7^M^−1^s^−1^) to generate faster assembly kinetics, with then the dissociation rate given by:

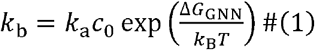

### 2.4 Testing self-assembly with NERDSS simulations

From the CG models, reaction networks, and reaction rates determined above, the final primary inputs to the NERDSS simulations are the subunit copy numbers and simulation parameters, including simulation volume. By default, we choose copy numbers by setting total monomers to 1*μ*M, split stoichiometrically across all subunit types. For fast simulations (minutes wall-clock), we specify 75 total subunits, and thus a simulation volume of (500nm)^3^. The time-step for simulations are determined automatically from the density (typically ∼0.1-1*μ*s) and 10^6^ iterations then produces a full second of assembly trajectory. By storing coordinates at fixed intervals in both .xyz and .PDB formats, our trajectories can be visualized with OVITO(65), and converted into Simularium files(64) for easy web-viewing and exploration.

For structure validation of target assembly from monomers, we also provide an alternate simulation initialization protocol. This protocol is designed to mimic a Lego-like build of the target complex so that it can be aligned and quantified relative to our designed CG structure. Our one-at-a-time titration protocol uses irreversible binding reactions to validate that each subunit can add together to form the target complex. Biopython routines then align this structure with the target structure to quantify the RMSD and report the agreement. We further test if all possible bonds are indeed present in the final structure using a NERDSS output file. We note that if a PDB structure represents only a fragment of a larger assembly, as in the HIV lattice, assembling an 18-mer of monomers through stochastic binding events may not align the final assembly with the complete target structure, although sub-structures should properly align. NERDSS supports several other simulation parameters such as filing writing frequency and a steric overlap distance check for subunit centers of mass following association events. We use NERDSS default values, noting these are readily edited in the sim_parms.inp file.

### 2.5. Non-spatial ODE models for comparing kinetics and equilibria

ioNERDSS constructs a corresponding non-spatial deterministic model for a given macromolecular complex for comparison with NERDSS trajectories. We call SciPy ODE solvers with a default stiff integrator (BDF) to integrate these deterministic and efficient mass-action rate equations (ODEs), compactly written as 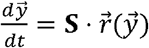, where 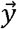 is the vector of species concentrations, **S** is the reaction stoichiometric matrix, and 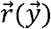 is the nonlinear vector of reaction propensities. However, we restrict the automatic construction of these systems of ODEs to complexes that contain <12 chains, due to the combinatorial growth in all possible intermediates and the need to enumerate only structurally admissible complexes (this can be over-ridden). The species 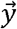 (monomers, intermediates, and the final complex) and reaction network for a given macromolecular complex is built from the coarse-grained graph of chains and edges (interactions) of section 2.1. We enumerate all subgraphs with an efficient method from recent work implementing Avis-Fukuda reverse search(70, 71). We note that for these models, like in recent work(9), we assume when a subunit binds to a complex, they form the maximal possible number of bonds immediately, to eliminate additional intermediates and a requirement for subsequent first-order reactions (SI). We do not restrict to monomer growth, allowing multi-subunit intermediates to bind together if sterically possible, just as in NERDSS. We define the rate constants to match the NERDSS model as closely as possible. The ODE models absorb the effects of diffusive travel time into macroscopic rate constants 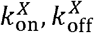, which differ from the intrinsic or microscopic rates 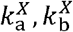, used in the NERDSS simulations. In 3D this conversion to a single macroscopic rate is well-established and accurate, 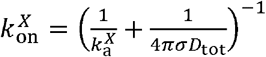, where *σ* is the binding radius of the reaction and *D*tot is the sum of the reaction partner’s diffusion coefficients (72). We include stoichiometric factors for binding between repeated subunits. Further, we note that the equilibrium in both approaches is expected to match (if the system is not kinetically trapped), because 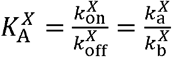 (72). Because association and dissociation events require all of a subunit’s bonds are made or broken per event, we specify that association rates between subunit *i* and a complex *j* that contain *m* bonds follows: 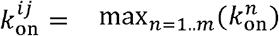. Because the ODEs only allow a monomer to associate through a single path (all bonds), whereas NERDSS allows each pair to form, the kinetics of higher order assembly will differ unless we account for these differences in pathways (see SI). The off-rates are then calculated based on dissociating all bonds present using:

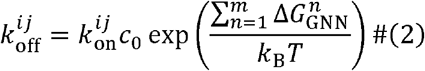

Note Eq. 2 thus differs from the purely pairwise specification for NERDSS in Eq. 1. Once a multi-valent subunit is bound to a complex, NERDSS automatically tests for available co-localized pairwise reactions, which are evaluated as first-order reactions(8).

## 3. Results

### 3.1. Automated generation of simulation-ready NERDSS models from PDB structures

We developed ioNERDSS as a user-friendly software tool which converts protein structure files into fully parameterized NERDSS models that are immediately ready for simulation (Figure 1). Given only a PDB identifier or file, ioNERDSS executes an automated workflow that extracts molecular geometry, identifies interaction interfaces and stoichiometry, estimates binding energetics, and produces all molecular and system files required for rigid-body stochastic reaction-diffusion simulations. The coarse-graining is very fast, typically only taking a few seconds. The prediction of binding affinities is the rate-limiting step, as it currently takes ∼0.5-2 minutes per interacting pair, which for very large complexes can add up to >1 hour. However, it should only need to be done once. ioNERDSS is provided open source, but users can bypass the source code and instead import it as a Python library (e.g. like NumPy), for easier function calls from existing code, including our provided tutorials. The tutorials provide step-by-step instructions and relatively straightforward customization options for key steps along the pipeline. Coarse-graining of the input structure yields a highly reduced but geometrically faithful representation of each distinct protein in the macromolecular complex as needed for its interactions. We leverage existing Python libraries when possible to enhance reproducibility and extensibility, including Biopython for sequence and structure analysis(66), and ProAffinity-GNN for predictions of binding affinities(41). In addition, we constructed a web interface and host a server at nerdssdemo.org to allow users to easily try out our tool from a given structure file or PDB ID. In addition to assembly topology and association/dissociation rates, self-assembly is highly sensitive to the copy numbers or concentration of each subunit. By default, we set these values to a total of 75 total subunit copies to support a full second of stochastic simulation in only 1-2 minutes of wall-clock time on a laptop. Copy numbers are specified as a simple labeled list in the parms.inp file and are thus easily re-defined. Variations in subunit abundances initially (22) or in time (4, 9) offer perhaps the easiest control knobs to push the system towards productive assembly of complete complexes, as we employ in our structure validation routines and illustrate below for the proteasome.

**Figure 1.**
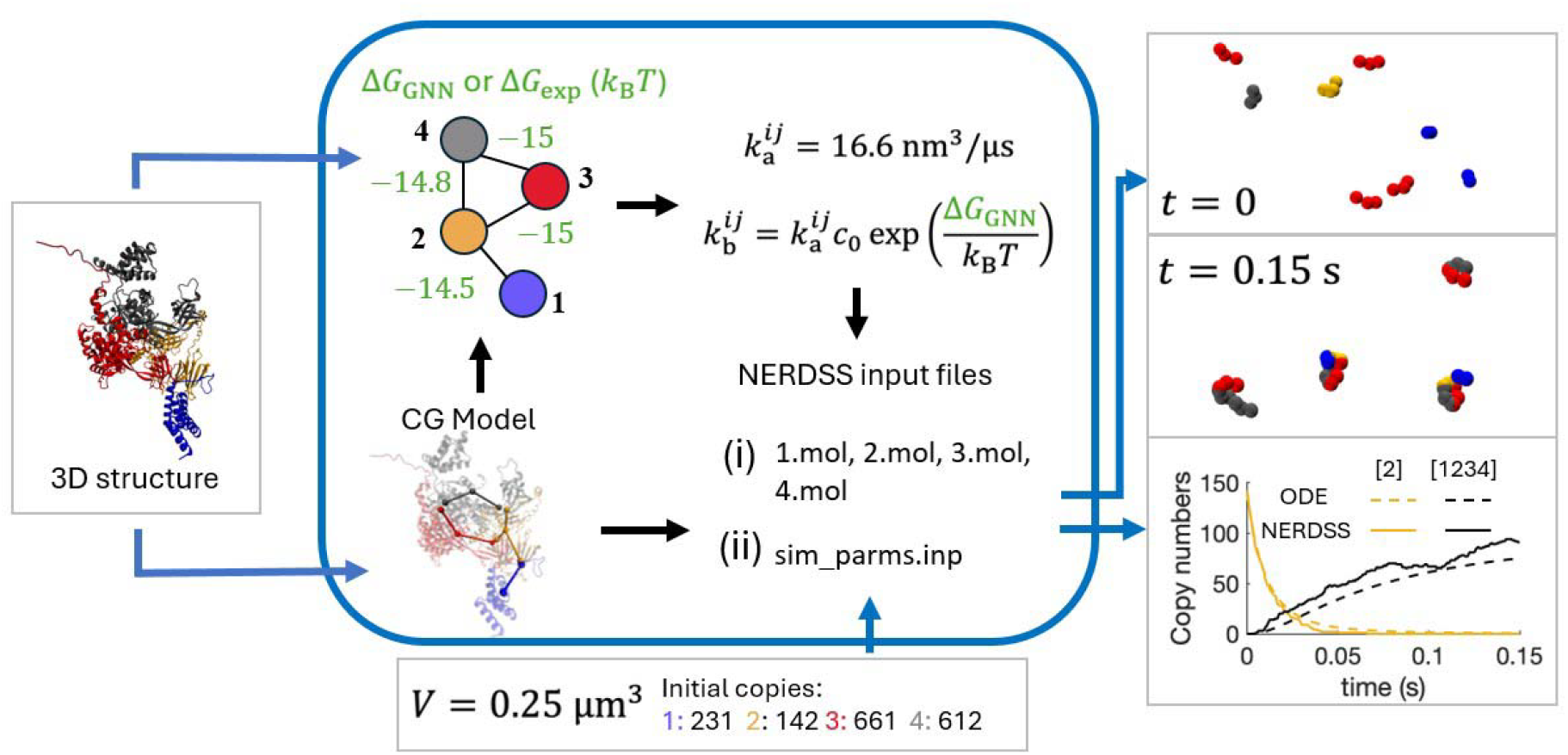
ioNERDSS converts atomic structures into executable models for self-assembly in NERDSS. Left column: the 4-subunit heteromeric R2TP co-chaperone complex was used as an example from ComplexPortal, where because the 3D structure has not been experimentally solved, we constructed it using AlphaFold3. Center: the blue box contains steps performed by ioNERDSS given the structure input on the left. The copy numbers/volume (lower box) are optional, with values used here from Saccharomyces Genome Database (SGD), scaled down by 20-fold for a smaller volume than the yeast cell. We otherwise use default values of 75 copies in total of all species so that simulating up to one second of self-assembly (Δ*t* = 0.25µs) completes in ∼ 1-2 minutes wall-clock time on a laptop, in a volume of 1.25 × 10^8^ nm^3^ (1µM). *c*_0_ = 1M and *i, j* are here indexing the subunits. Rates are for NERDSS simulations (for ODEs see Methods). Δ*G* is from ProAffinity. Right column: outputs include the coordinates of all subunits in the volume in PDB and xyz formats (top right), and time-resolved copies of all intermediates, with a representative monomer and the final tetramer shown (bottom right).

### 3.2. Constructing models from complexes with repeated subunits is idiosyncratic

When all complex subunits are distinct, the coarse-graining procedure is straightforward. Each chain-chain contact must have a distinct interface to mediate this new contact, and thus the total number of edges *N*_edge_ produces 2*N*_edge_ interfaces. Accurately coarse-graining protein complexes that contain repeated subunits is more technically demanding for automated model construction, yet it is critically important, as repeated subunits emerge commonly in multi-subunit complexes via evolution(73). We therefore introduced several additional criteria to distinguish homotypic vs heterotypic interactions between homodimers (Fig 2) and to accommodate variability in geometry or the molecular neighborhoods of nominally identical subunits (Fig 2B). For example, in the homotetrameric complex of Fig 2A (74), all four repeated subunits interact with all other subunits, producing *N*_edge_ =6 interactions. The minimum number of interfaces is 3, and ultimately we derive that there are only 3 distinct interfaces per subunit, with each engaging in a homotypic (e.g. yellow-to-yellow site) interaction. To classify the interfaces as the same type, we use geometric signatures of the CG interfaces (distance and orientation) and verify that the residues that comprise the interface are the same (Methods). Because of thermal variations in structures, we use thresholds to assess similarity (see Table S1).

**Figure 2:**
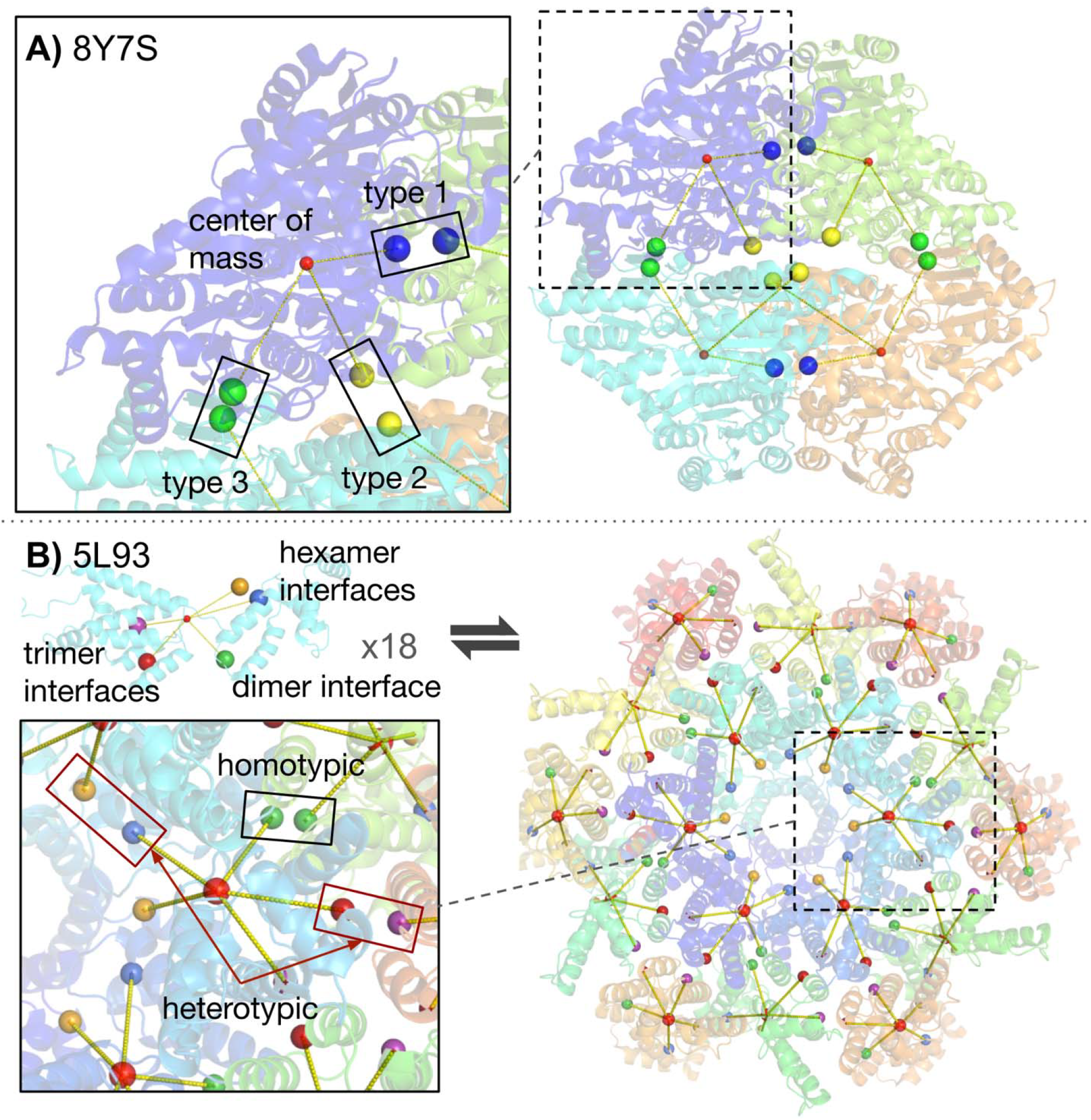
Macromolecular complexes with repeated subunits and variable stoichiometry or interface symmetries are more technically difficult to coarse-grain. A) Ribbon structure and ioNERDSS coarse-grained structure of a benzaldehyde lyase mutant M6 from PDB ID: 8Y7S. 8Y7S is an example of a homo-tetramer that therefore coarsens to a single subunit, A.mol. Each monomer binds with the three others using a distinct homotypic interface, yellow-to-yellow, blue-to-blue, and green-to-green, with the COM in red. B) Ribbon structure and ioNERDSS coarse-grained structure of a fragment of the HIV-1 immature Gag lattice from PDB ID: 5L93. 5L93 also coarsens to a single subunit, Gag.mol, but each subunit contains both homotypic contacts stabilizing a dimer interface (green-to-green) and heterotypic interactions that stabilize hexamers (blue-to-orange) and trimers (red-to-purple), supporting the continual growth of a much larger lattice than the visualized fragment. The outer monomers appear to be in a distinct chemical environment, but by recognizing the sequences as the same protein, ioNERDSS identifies the maximum number of interfaces (here 5) and projects a single rigid template onto all other repeated chain positions. For structures like the immature lattice that live on a spherical shell, we can optionally improve the regularization of the subunit positions to ensure a single spherical curvature is achieved during stochastic assembly.

For the HIV immature Gag lattice example (Fig 2B), the resolved structure contains one subunit repeated 18 times, but as a truncated part of a larger assembly containing thousands of repeats (3). As a result, the monomer subunits are in three different binding states, bound to either 5 partners, 4 partners, or 3 partners. Here again we rely on our geometric and residue-level criteria to identify one homotypic interaction and two heterotypic interactions that are then mediated by a maximal set of 5 interfaces per subunit (Fig 2B). A challenge with this large homomeric lattice is that because it can be built via multiple distinct pathways of interface contacts (dimer, or hexamer, or trimer first, in principle), small local alignment variations when averaged over repeated subunits can give rise to global alignment issues, such as distinct curvatures when following dimer vs hexamer growth pathways. This variability may also arise because this tri-hexagonal lattice cannot perfectly tile a sphere and thus must embed defects in lattice structure. However, to enforce a single curvature of growth, we can exploit symmetries in the known structure. By constraining each rigid subunit to also lie on the surface of a sphere of radius *k*, we can ensure the target lattice is formed with a single curvature of 1/*R* and with growth to a complete (with defects) spherical lattice containing >3000 subunits (4, 67). This also means we can design the same lattice architecture to adopt a higher curvature, for example, producing smaller spheres with fewer subunits required. This option is available for any spherically ordered complex structure.

There are exceptions where a subunit truly does inhabit a distinct molecular neighborhood than its repeated copy, such as the human GATOR2 complex (PDB 7UHY) (75). The assembled structure contains two repeats of the MIOS subunit, but one repeat binds a partner WDR59, whereas the other binds a different partner WDR24. These interactions use distinct residues and contact orientations within the assembly, requiring distinct interfaces. However, they are also mutually exclusive, such that one MIOS subunit could not be bound to both partners simultaneously. Because NERDSS supports conditional interactions, in this case we can assign reaction rules where either binding interaction is only possible if the other interface is still unbound, preventing steric overlap during assembly. However, because this is less common than minor local variations in interface sites, we implement explicit user warnings when repeated subunits do not produce consistent interface sets. These examples illustrate that special care can be required to ensure inferred interfaces are complete and correct if repeated subunits are present. The next section systematically benchmarks this process and provides tutorials to validate and troubleshoot assembled CG models.

### 3.3 Structures are validated using fast assembly protocols

We provide routines and tutorials to validate that the CG structures that will form via stochastic assembly will align with the designed target structure. First, however, we verify that all *N*_chains_ identified in the PDB file are connected into the single graph representing the target structure. If instead, some subunits are disconnected from the graph, the subunits cannot possibly assemble the target *N*-mer, and users are instructed to increase interface detection thresholds. Given a single connected graph, we then test stochastic assembly. Instead of running a NERDSS simulation with reversible interactions at 1 μM concentration, the validation protocol is designed for fast and irreversible association of subunits at higher concentrations. To accelerate self-assembly, we apply higher concentrations via a smaller simulation box (50 nm side length square box), faster association rates, and no dissociation (set to 0), which will drive fast bimolecular association, but will also easily lead to kinetic traps(9). To avoid kinetic traps, we initialize the simulations with subunits enough to assemble only one target structure. We identify this structure from the simulation trajectory, align the coordinates of this structure using rigid-body transformations with the target designed structure, and report the RMSD between the COMs. In addition to baseline benchmarking (SI), to support validation of structures with repeated subunits, the protocol can also support titration of additional subunits, which when done sufficiently slowly ensures that kinetic traps are minimized and each new subunit adds onto existing nucleated structures(4).

We ran validation on the >44,000 PDB structures that contain 3 to 12 protein chains, restricted to protein-only entries, experimental methods (X-ray diffraction or electron microscopy), exact protein chain count (protein entity count and number of chains matches in the PDB entry), resolution (< 3.5 Å). We skipped dimers because they are straightforward to coarse-grain. Fully heteromeric structures were reliably assembled to the target structure with RMSD < 10^-6^ nm, indicating excellent alignment. For these structures, stochastic assembly was also quite reliable, with only <1% failing to stochastically assemble to the final structure (termed ‘under-assembly’). However, 40% of these structures were not simulated because they did not produce singly connected graphs; the structures or hyperparameters must be interrogated more closely to assess missing interfaces in these cases. For structures with repeated subunits, only 22.5% contain disconnected graphs and 3% were excluded for identifying <2 chains in the target. Of the remaining 75%, 1.4% failed to stochastically assemble to the final structure (under-assembly). The remaining 73% assembled to a complete target structure, although here we did see higher RMSD between the designed structures COMs and the stochastically assembled COM. A majority (61% of the successful ones) still had a very accurate RMSD of <0.1 nm, while the remaining structures produced higher RMSD that could be perfectly acceptable or indicate an inaccurate target.

To troubleshoot structures that i) produced disconnected graphs, or ii) during simulation did not complete assembly over the stochastic simulations (under-assembly) or iii) did complete assembly but to a target structure with a larger RMSD that might be inaccurate, we provide a Python tutorial that is designed to test these three primary failure modes. In the simplest case, a disconnected graph indicates one or more subunits do not bind to the complex and increasing *N*_cut_ and *r*_cut_ provide a more permissive threshold to identify binding interfaces between subunits. For the case of under-assembly, no assemblies were found that reached the target size, and the issue could simply be that the simulations need to be run longer, particularly for large homomeric structures, so subunits find the remaining open binding interfaces. Finally, even if the target structure is found, if the structure has repeated subunits, the RMSD can be too high. The most likely culprit are structure issues with the CG subunits. Structures that identify too many interfaces are most likely to create assembly errors and amorphous, poorly structured assemblies. This can occur with repeated subunits if functionally identical interfaces are instead assigned to be distinct. Decreasing *N*_cut_ and *r*_cut_ thresholds can then improve this and define fewer CG interfaces. Even if the interface contacts look correct, a less common issue due to imperfect alignment within structures can be multiple copies of one subunit stochastically binding into a location where only one is expected, which occurs for the proteasome example below. NERDSS uses a simulation parameter (overlapSepLimit) to checks for steric overlap between each subunit that associates into a complex by measuring the distance between the subunit COMs. This threshold is typically set quite low (0.1nm), as most assembled structures do not contain ‘defects’ or imperfectly aligned repeated subunits. By increasing this threshold, we can ensure two subunits can never occupy the position of a single CG subunit, and we report the appropriate upper bound to ensure correctly bound partners will not be rejected due to steric overlap. Therefore, for homomeric structures or partial repeated subunits, titrating in additional subunits is an important final validation step, to ensure simulations do not produce ‘over-assembly’, where a finite-sized structure like the proteasome with 28 subunits continues to add more subunits in a spiral-like growth. A final issue is that a PDB of a partial homomeric assembly (like the 18-mer HIV Gag lattice or the 8-mer actin assembly) represents a specific geometry of repeated subunits. Stochastic simulations may produce a target *n*-mer with the correct number of subunits but not assembled into the exact same geometry, which will produce differences in the RMSD, even if the subunits are adopting the correct topology. In this case the CG model and the simulations are correct—it is rather that statistically the PDB structure simply represents only one of many acceptable *n*-mers. We note that having too few interfaces within the complex can lower the stability of the structure and will change accessible assembly pathways, but it does not typically lead to assembly errors. If all of our proposed strategies for correcting the assembly fail, our Python tutorial directs users to raise a GitHub issue so we can assess the system more deeply.

### 3.4. ioNERDSS launches NERDSS simulations with matching ODE models for smaller complexes

Once the topology of the macromolecular assembly has been deconstructed by ioNERDSS into its constituent subunits, we can study assembly pathways. The simplest initial conditions are to randomly distribute all subunits as monomers throughout the simulation volume. Specifying copy numbers (concentrations) to known values, such as reported by cell-type, is straightforward through our text-based input files. By default, we choose copy numbers by setting total monomers to 1uM, split stoichiometrically across all subunit types. With estimated binding affinities from ProAffinity-GNN typically in the nM-to high uM regime, this concentration is often high enough for higher-order assemblies with stabilizing loops to form (Fig 3). By choosing relatively fast association rates and only ∼hundred subunits, assembly proceeds quickly to allow for rapid assessment of structures formed within a few minutes wall-clock time. ioNERDSS provides simple analysis routines in its Analyzer module that read in the formatted outputs generated by the NERDSS simulations to quantify yield of any specie(s) vs time (Fig 3).

**Figure 3.**
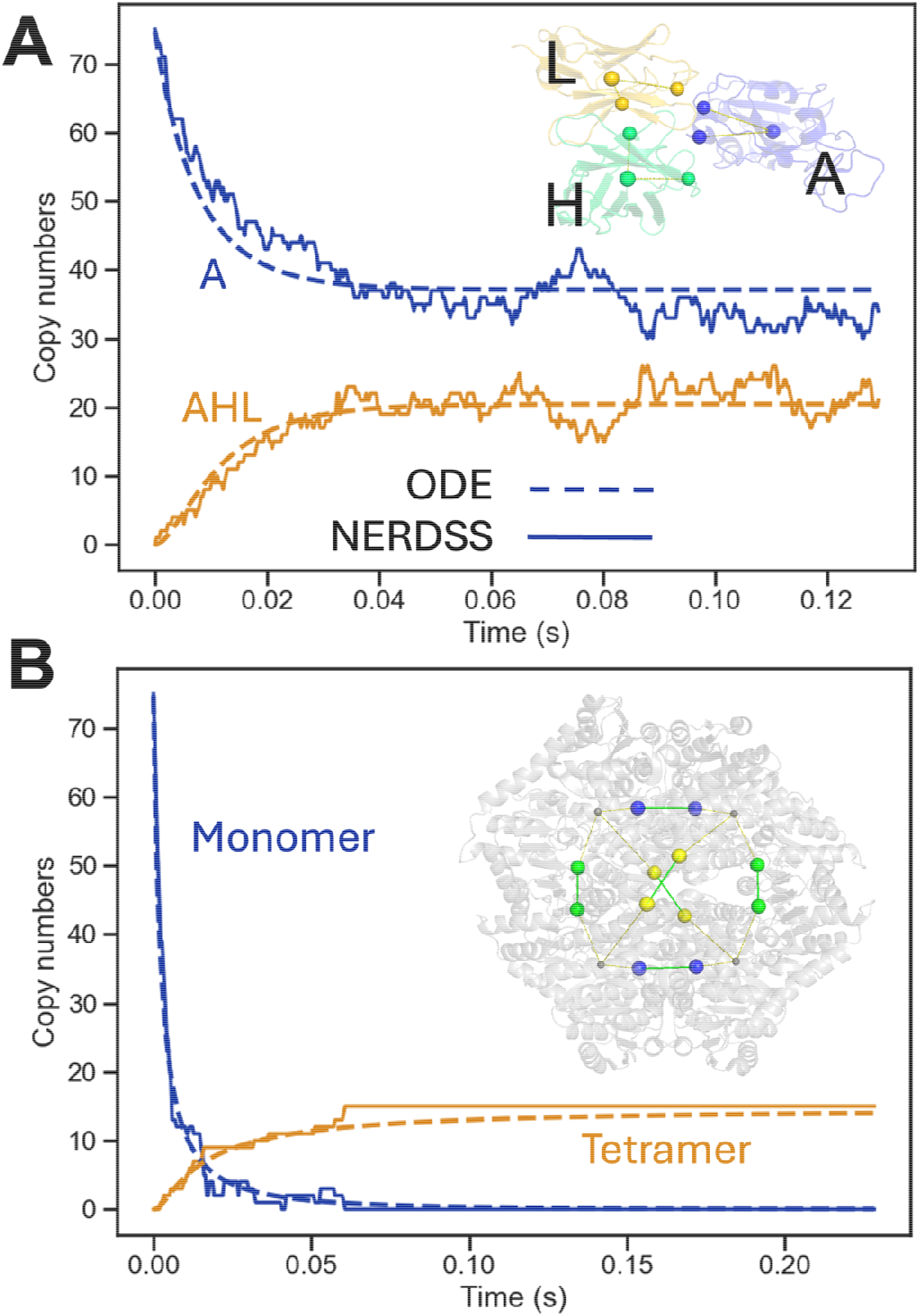
ioNERDSS compares assembly kinetics for stochastic NERDSS assembly with generated ODEs for small complexes. For these NERDSS simulations of relatively small assemblies with randomized, well-mixed initial conditions, we expect and observe strong agreement to the ioNERDSS-generated ODE models. A) For heterotrimer 8ERQ, each subunit is distinct and interacts with both partners, forming a closed cycle. Initial copies for each subunit are 130 in a volume of (600 nm)^3^, and we plot Chain A (blue) and the total trimers (orange). One NERDSS trajectory in solid lines, and ODE in dashed. B) Homotetramer 8Y7S was initialized with 130 monomer copies in a simulation box of (600 nm)^3^, monomer in blue, completed tetramers in orange. Inset shows the PDB structure in gray, and the CG monomers with blue, yellow and green interfaces, and gray COM. Green lines indicate intermolecular contacts. NERDSS simulations were run with a time step of 0.01 μs.

With smaller complexes (<12 subunits), ioNERDSS enumerates all intermediates to construct a complete set of rate equations to track species vs time deterministically and in the well-mixed approximation, taking care of stoichiometric factors for repeated subunits and assigning rates that will maximize agreement with NERDSS (see Methods). These systems of ODEs are numerically very efficient to integrate and in Figure 3 we compare the assembly kinetics using the same initial conditions and rates to the NERDSS simulations of a homotetramer (four total species) and a heterotrimer (7 total species: 3 monomers, 3 dimers, and 1 trimer). We see very good agreement as expected, given that the NERDSS simulations are also ‘well-mixed’ initially, the assembly is all occurring in 3D, and we incorporate the impact of diffusion into the macroscopic rate constants (76). The rate equations further allow non-monomer growth pathways (e.g. dimer + dimer), just as in the NERDSS simulations. The similar kinetics emerges in both the spatial stochastic simulations and the ODEs despite differences in the allowed intermediates; in the spatial simulations, only pairwise contacts can form per timestep. For example, the addition of a monomer to a dimer will form only a single bond leading to an ‘open’ trimer, whereas in the ODE models, we specify that the addition of the monomer will complete the closed trimer. However, in NERDSS, the ‘open’ trimer now has a co-localized pair of interfaces that can bind following a first-order reaction (they are not diffusing relative to one another), which typically occurs significantly more rapidly than dissociation, leading to a closed trimer. The lifetimes of the combined trimer states (open and closed) thus agrees well with the lifetime of the single closed trimer specie in the ODE model, which is controlled by Eq. 2. With the larger complexes in the next sections, the combinatorial growth in species renders the deterministic ODE approach extremely costly, whereas the specification of the NERDSS model is still of the order *N*_edge_ .

### 3.5. Large assemblies and filaments are tractably defined and simulated with ioNERDSS

The pseudo-symmetric 28-subunit proteasome exemplifies several of the distinct features handled via the ioNERDSS coarse-graining and simulation pipeline. It contains 14 distinct subunits (two 7-mer rings), each repeated twice, with 48 interactions, all of which are heterotypic except two (Fig 4A). ioNERDSS successfully identifies the interfaces and interactions, although due to a fair amount of structural heterogeneity within the contacting 7-mer rings, we adjusted the default hyperparameters (SI). Specifically, we had to increase the *r*_cut_ to make sure all interfaces were detected. Secondly, we had to increase a stochastic simulation parameter that checks for steric overlap to be more permissive, so two proteins would not be able to occupy the site of a single binding partner (SI). The binding affinities for all contacts are predicted to be relatively similar in strength, resulting in a highly connected and stable loop-forming assembly topology that will easily suffer kinetic traps(9). Because we initialize all models with the maximally permissible set of interactions by default, such that contacts in all rings can form immediately, this lack of built-in hierarchy starves the system of monomers. Several protocols could help resolve this, and biologically, the proteasome is known to assemble in two equivalent halves prior to slow dimerization of these halves, enforcing a hierarchy of timescales (77). To demonstrate the completed 28-mer assembly (Fig 4B,C), here instead we use a simple but effective protocol of titrating in monomers slowly enough (*k*_titr_ = 2 − 9 × 10^-6^ M/s) that early nucleation events are succeeded by elongation and completion rather than competition from many other nucleation events (4, 9).

**Figure 4.**
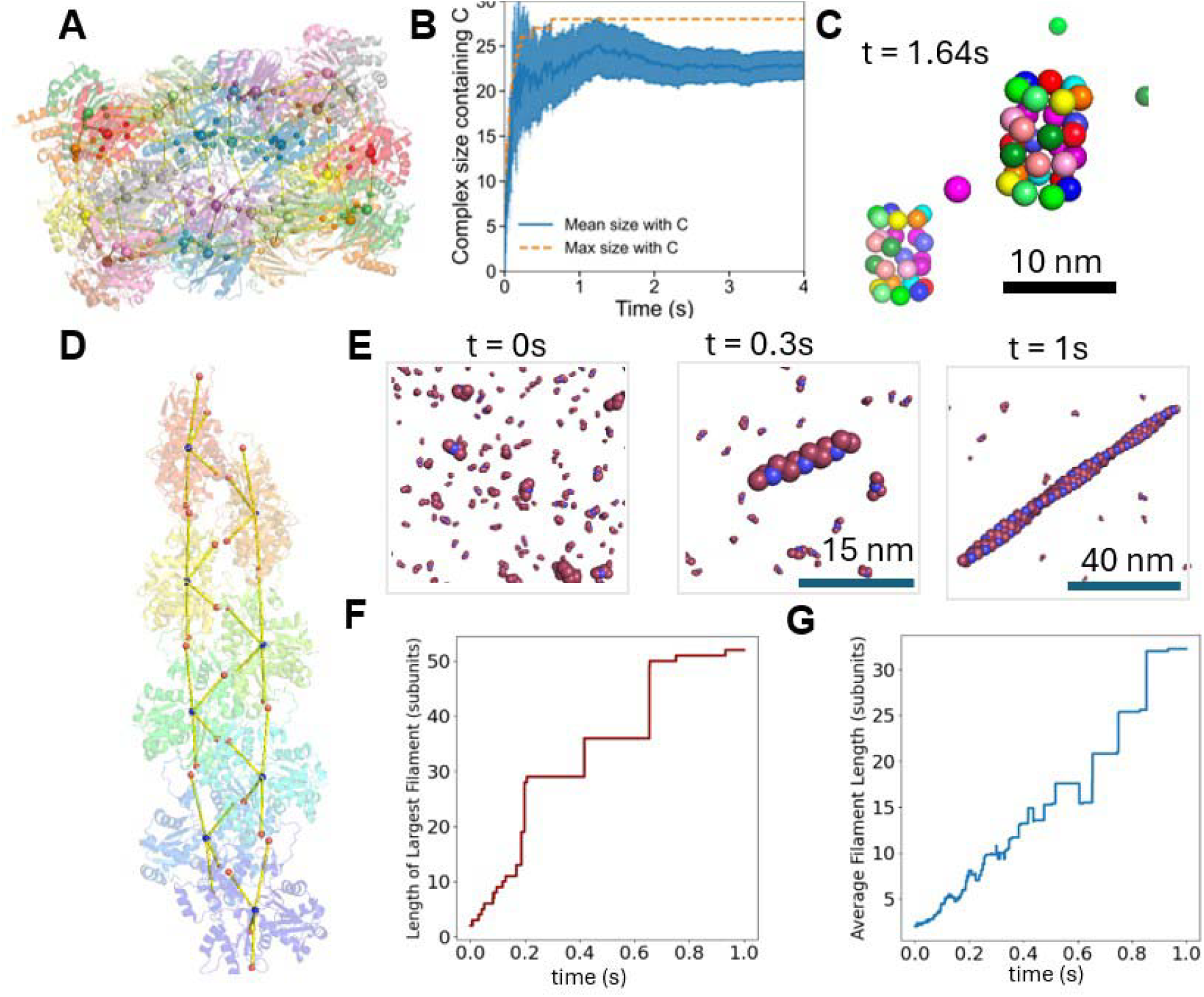
ioNERDSS resolves CG monomers of multi-component machines and single-component filaments. A) Proteasome is constructed of 14 distinct subunits, with each appearing twice. B) Mean and max complex size containing proteosome chain C over time shows that slow titration of monomer subunits in the simulation volume supports assembly of the complete 28-mers, along with other intermediates. Error bars show +/- 1 standard deviation. Mean complex size and standard deviation is calculated from one trajectory averaging the number of subunits of all complexes with C in each time step. C) Snapshot of the completed 28-mer from a NERDSS simulation. Each color represents a distinct type of subunit. D) Actin monomers (6BNO) assemble into filaments. The ioNERDSS CG model of actin monomers derive four distinct interfaces per subunit (red circles) relative to the subunit center-of mass (blue circles). The four interfaces (top, bottom, cross up, cross down) participate in two heterotypic interactions total, top-to-bottom and cross up-to-cross down. The positions of interface sites and orientations between bound monomers stabilize formation of the known helical actin filament. E) Snapshots of the NERDSS simulation starting from 130 monomer copies in a 600 nm side-length box (1 monomers). F) ioNERDSS analysis tools quantify the length of the largest filament vs time and G) the average filament length vs time, shown here in number of subunits.

Actin filaments represent a canonical self-assembly system whose pseudo-one-dimensional growth has no inherent geometric or structural termination point. Using a high-resolution structure of 8 actin monomers assembled into the F-actin polymer (78) (6BNO), ioNERDSS identified four distinct interfaces per monomer (Figure 4A), which together support the characteristic helical twist of the filament. Simulated assembly trajectories with NERDSS at 1*μ*M of monomers with *K*_D_ values of 0.01 *μ*M on the long axis and 2,000 *μ*M on the cross-axis produce nucleation and growth of long >120 subunit polymers (Fig 4C). This persistent growth beyond the 8 monomer PDB structure size demonstrates how the subunits and contacts are correctly learned via the ioNERDSS coarse-graining procedure to generate structurally faithful assembly pathways.

### 3.6. Ideal platonic solids offer design templates for self-assembly

The five Platonic solids are ideal assembly structures with each built of identical subunits and orientational contacts: the tetrahedron, cube, octahedron, dodecahedron, and icosahedron. The platonic routines in ioNERDSS provide functions to generate each of the five platonic solids without any additional input and with easily tunable sizes. In Figure 5 we illustrate the results from simulating dodecahedron subunits over time, ultimately generating multiple fully assembled 12-mers. Subunit centers are placed at face centers, and interfaces are placed at each edge of a Platonic solid, thus each 12-mer subunit has 5 interfaces. To control the size of the solid, users input the circumscribing radius of the solid. Users can also increase the binding radius *σ* to increase the excluded volume of the individual interfaces. The icosahedron is a common topology for viral capsids like Hepatitis B virus (47, 48), and with this library, users can bypass construction using imperfect PDB structures to create tunable CG subunits for icosahedral assembly.

**Figure 5:**
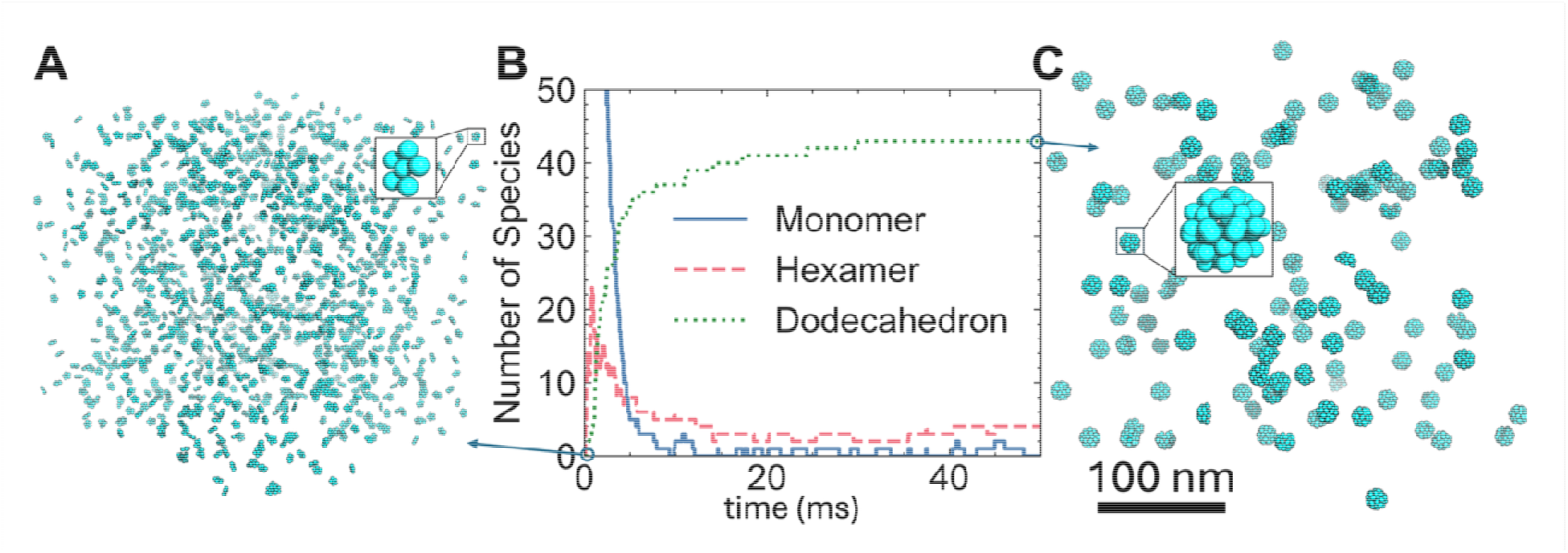
Simulating assembly of the tunable dodecahedron Platonic solid. Ideal structures like dodecahedrons have a fixed geometry and the Platonic library of ioNERDSS designs the monomer structure and interactions to ensure a target radius of the completed assembly is achieved. (A) Here, so each subunit (inset) has 5 interfaces separated from the center of mass by 4.8 nm. Snapshot at time . The simulation was run at 50 of monomers with per pairwise interaction. (B) All subunits are initialized as monomers. We show here the species evolution of monomers (blue), hexamers (red dashed), and completed dodecahedrons (green dot) in time. (C) After 50 ms, monomers have all assembled into completed dodecahedron (inset) or incomplete intermediates.

## 4. Discussion

Defining biological models for simulations using reaction-diffusion software like NERDSS is notoriously not automated, relying on the user to decide which components to include and which interactions or reactions they obey. While this stems from the broad flexibility of rate-based modeling approaches, it creates a barrier for non-experts to explore and quantify the dynamics of mesoscale processes like macromolecular self-assembly subjected to thermodynamic constraints. With ioNERDSS, we therefore narrow down the possible reaction networks that one can dream up in tools like Catalyst(79), to build a thermodynamically consistent reaction network directly from 3D atomic structures. This removes a substantial barrier to starting stochastic RD simulations by automating the set-up of simulation-ready models and supports orthogonal validation strategies of equilibrium steady-states. We do so by translating atomic structures into rigid-body coarse-grained models with accurate interface geometry and appropriate reaction rules — a process that previously required substantial manual intervention. With immediate applications to characterizing mechanisms and control of multi-valent protein self-assembly, these models then provide a starting point to expand to even richer cell-scale systems, including the addition of membranes(38, 39, 80), receptors(81), enzymes(76, 82), and nucleic acids (83). These models also support assembly of disordered systems, by simply removing orientational constraints between subunits to allow diverse contact angles between bound rigid subunits. Automating the model instantiation process, much as software tools have done for MD simulations(42), helps improve accessibility, reproducibility(62), and expansion of these tools for mesoscale out to whole-cell models.

By building ioNERDSS as an accessible library within the Python ecosystem, we can immediately and in the future integrate readily with state-of-the-art machine-learning and AI-based tools for model optimization and analysis. ioNERDSS currently incorporates structure-derived predicted binding affinities(41) into our reaction-networks by converting them into association/dissociation rates, making it possible to predict emergent assembly behavior from local, structure-based interactions. A fully ‘bottom-up’ approach to predicting binding rates between all subunits and intermediates in the assembly pipeline is not readily tractable with MD simulations without substantial expertise(55), but tool development would make this approach a natural future step in the ioNERDSS pipeline. A more immediate pathway to learn the kinetics of the assembly pathway is via a ‘top-down’ or data-driven approach, from experimental data. While this requires time-resolved assembly kinetics experiments, the availability of a ready-to-simulate assembly model from ioNERDSS is compatible with various optimization strategies to learn model parameters from data(9). Because NERDSS offers the flexibility to encode allosteric effects due to conformational changes, for example, in reaction rules, it does not require that proteins were in rigid conformations throughout assembly. This supports exploration of an even larger space of assembly pathways, where binding between partners changes conformations to significantly alter the kinetics of subsequent interactions(84).

ioNERDSS is designed to systematically handle repeated subunits to ensure target topologies are realized upon stochastic simulation, but there are limits due to the rigid-body assumptions of the NERDSS simulator. The clathrin cage and the HIV mature lattice are two single-component assemblies that contain both hexameric and pentameric cycles in the completed structures. Unlike the ideal structures of the Platonic solids, these topologies require varying binding angles between otherwise identical subunits. Because ioNERDSS will assign a single set of interfaces and bond orientations, simulated assemblies will then contain defects instead of the experimentally observed closed pentagons. This can also present a challenge for ultra CG MD, where relatively rigid subunits are also the default. The addition of energy-function based interactions between subunits is needed to robustly correct for these topological variations while retaining rigid subunits(85). Clathrin trimers have a further complication; what we think of as the assembly subunits, the clathrin trimers, are actually hexamers of 3 light and 3 heavy chains. Our current NERDSS trimer models were thus designed by hand(38, 81), as ioNERDSS identifies light chains and heavy chains as the assembly subunits. A natural extension is to allow users to merge chains to create larger subunits for self-assembly. Further, because non-protein components like nucleic acids(83) and membranes(38, 39, 80) often play essential roles in self-assembly, automating the addition of these structures with contacts to protein subunits represents an important extension.

To conclude, the process of self-assembly is an essential and in many cases highly controlled step in homeostasis(10) and decision-making by living systems(81). The simulation-ready models from ioNERDSS for any 3D structure offer a highly efficient method to explore and optimize models relative to experimental datasets, offering a foundation for broad and quantitative studies of self-assembly pathways and control mechanisms with hundreds or thousands of subunit copies. It also offers a method for kinetic design, as selection of binding rates or kinetic protocols that assemble high yield in realistic time scales can be a challenging problem(9). Recent work on multi-subunit, semi-addressable self-assembly demonstrates that by carefully designing binding interactions, one can optimize both assembly yield and efficiency while suppressing off-pathway structures (23, 25). ioNERDSS connects these approaches to assemblies built from protein subunits, assembling diverse 3D topologies. Our automatically constructed reaction networks enforce thermodynamics of higher-order reversible assembly, helping reveal intrinsic thermodynamic or kinetic barriers to robust assembly. As structural databases continue to grow rapidly, and as binding-affinity prediction improves, ioNERDSS provides a practical, reproducible, and transparent framework for hypothesis testing, model sharing, and broad community use, available open-source at github.com/JohnsonBiophysicsLab/ionerdss.

## ACKNOWLEDGMENTS

MEJ gratefully acknowledges funding support from US National Institutes of Health research awards R35GM133644 and NIAID R01AI186663 supporting MEJ, YMY, MS, JAF, SLF and SG. SLF also gratefully acknowledges postdoctoral funding support from the Gordon and Betty Moore Foundation.

