## Supplemental Information for "Transforming macromolecular structures into simulations of self-assembly with ioNERDSS"

for

### SUPPORTING TEXT AND METHODS

#### 1. Implementation details for the PDB to NERDSS parameter pipeline

##### 1.1. Parsing the structure for coarse-graining

We convert all-atom protein complexes (from mmCIF / PDB file) into a coarse-grained representation suitable for NERDSS reaction-diffusion simulations by determining interfaces between polypeptide chain pairs via KD-tree queries over Cα positions with a distance cutoff. Each validated interface is summarized by the mean of contacting Cα coordinates, a set of contacting residue identifiers on both partners.

Atomic coordinates were parsed from mmCIF / PDB using the Biopython package. We identify $N_{\text{chain}}$ polypeptide chains in the file, where only chains containing at least one standard amino acid are retained, and chain identifiers are sorted to ensure deterministic ordering. Throughout, positions are treated in angstroms (Å) sticking to the units from mmCIF / PDB, while the user-facing hyperparameters such as distance cutoff are specified in nm.

For each chain, a center of mass (COM) and coarse-grained (CG) radius are computed from all atoms in standard amino-acid residues present in the given mmCIF / PDB file. If hydrogen atoms are present in the provided mmCIF / PDB file, they are also considered. We do not add or remove hydrogen atoms in this step. Let chain $C$ consist of atoms with Cartesian coordinates $\vec{x}_a$, where $a = 1 \dots N_C$ and $N_C$ is the total number of atoms on chain $C$. The chain COM $\vec{c}$ is defined purely geometrically (unweighted by mass):

$$\vec{c} = \frac{1}{N_C}\sum_{a=1}^{N_C} \vec{x}_a \tag{M1}$$

Similarly, the CG radius $R_C$ is also unweighted by mass, defined as root-mean-square distance of atoms from the chain's geometric center (centroid):

$$R_C = \sqrt{\frac{1}{N_C}\sum_{a=1}^{N_C} \|\vec{x}_a - \vec{c}\|^2} \tag{M2}$$

The chain COM and CG radius are used to represent the position and estimated size of the chain downstream.

##### *1.2. Interface determination between chains in the complex*

To avoid iteration over distances between all chain pairs, we can avoid checking across residues of distant chains that are very unlikely to interact. We first compute axis-aligned bounding boxes for each chain from all atomic coordinates. Two chains $C_i$ and $C_j$ are considered too distant to interact if their axis-aligned bounding boxes are separated by more than $r_{\text{cut}}$ (the interacting residue distance cutoff) along any axis. Formally, let $(m_{k,d}, M_{k,d})$ denote the minima and maxima of the axis-aligned bounding box of chain $k$ along axis $d$, then the pair of chains $C_i, C_j$ is rejected as interacting if:

$$\exists d \in \{x, y, z\} \left(m_{j,d} - M_{i,d} > r_{\text{cut}} \text{ or } m_{i,d} - M_{j,d} > r_{\text{cut}}\right) \tag{M3}$$

We assume that for any two chains $C_i, C_j$, there can only be either zero or one interface between the two chains. In other words, if multiple residue patches mediate contact between two chains, they are coarse-grained to one effective interface. We determine that a pair of chains $\left(C_i, C_j\right)$ is **interacting** if each chain contributes at least $N_{\mathrm{cut}} = 3$ (residue cutoff) residues that lie within $r_{\mathrm{cut}} = 6A$ (distance cutoff) of residues on the other chain. For a candidate pair $\left(C_i, C_j\right)$, we build a KD-tree on the Cα coordinates of $C_j$ and query neighbors for each residue in $C_i$ within radius $r_{\mathrm{cut}}$. The number of Cα in $C_i$ that are in contact with $C_j$ is denoted as $N_{ij}$. We also count $N_{ji}$ for the number of Cα in $C_j$ in contact with $C_i$. If both $N_{ij}$ and $N_{ji}$ are greater or equal to $N_{\mathrm{cut}}$, the pair $\left(C_i, C_j\right)$ is flagged as having an interface; the interface point for each chain is taken as the geometric mean of its contacting Cα coordinates. If either interface is too small ( $N_{ij} < N_{\mathrm{cut}}$ or $N_{ji} < N_{\mathrm{cut}}$), then we do not assign an interface between these two chains.

Interface detection is enforced to be symmetric: a validated contact between $C_i$ and $C_j$ produces entries on both chains, including an interface on $C_i$ that binds to $C_j$ and an interface on $C_j$ that binds to $C_i$. We maintain an explicit binding partner index map $\pi_{\mathrm{intf}}$ to bookkeep the pairwise relationship of interfaces.

$$\pi_{\mathrm{intf}}: (i, a) \mapsto (j, b) \tag{M4}$$

where $i, a$ refers to the $a$-th partner on chain $i$ and $j, b$ refers to the $b$-th partner on chain $j$.

##### 1.3. Detecting repeated proteins

We first partition protein chains into equivalence classes of repeated copies of the same protein, resulting in a total number of distinct subunits $N_{\text{mol}}$ , where $N_{\text{mol}} \leq N_{\text{chain}}$ . For example, the HIV Gag lattice contains 18 chains but only $N_{\text{mol}} = 1$ protein, Gag. We do so by using metadata from the mmCIF header or similarity thresholds at the sequence level. We note that structural homology between distinct proteins is not truly relevant to the coarse-graining, as each distinct subunit has independent tunable copy numbers, even if binding affinities or rates are similar. The output is a map from each chain ID to a group representative and a list of chains per group, which we use throughout the pipeline to assign consistent molecule-template names and to reuse interface templates across repeated subunits. The parameter `matching_mode` switches between the three modes as in Table SB: header/read, sequence, structure. Header-based grouping is fastest and aligns with the depositor's intended biological assembly but is not always present or accurate in derived structures. If available, we read `_entity_poly.pdbx_strand_id` from the mmCIF header to obtain the set of chain IDs that belong to each distinct protein type. If the header field is absent or incomplete, we fall back to sequence-based grouping. Alternatively, we suggest the user provide strand ids in the input mmCIF / PDB, if the user knows the species of each chain. Thresholds $\delta_{\text{seq}}$ or $\delta_{\text{RMSD}}$ and the matching mode can be tuned through optional arguments or `PDBModelHyperparameters` to balance sensitivity and specificity for the system under study. The $\delta_{\text{RMSD}}$ is not recommended as it is structure based, and distinct proteins with low sequence homology can still have very high structural homology.

For sequence-based repeat determination, we compute amino-acid sequences with `Bio.PDB.Polypeptide.PPBuilder`, which extracts polypeptide chains as contiguous peptides. For chains $C_i$ and $C_j$ with sequences $s_i, s_j$, we perform polypeptide sequence alignment with a `Bio.Align.PairwiseAligner` and yield the alignment score $\text{align\_score}(s_i, s_j)$. We define identity score $\text{IDS}(i, j)$ as:

$$\text{IDS}(i, j) = \frac{\text{align\_score}(s_i, s_j)}{\max\{|s_i|, |s_j|\}} \tag{M5}$$

All $C_j$ satisfying $\text{IDS}(i, j) \geq \delta_{\text{seq}}$ (where $\delta_{\text{seq}} = 0.5$ is the sequence threshold hyperparameter default) are grouped with the current $C_i$. We update a visited set and continue until all chains have been assigned. This produces disjoint groups, each labeled by the first chain encountered.

##### 1.4. Molecule types and diffusion parameters

From the finalized molecular templates, we create the molecule type files used by the NERDSS simulator. Each molecule type contains the interface names and offsets copied from its template, together with translational ($\mathbf{D}_{\text{t}}(D_{\text{t},x}, D_{\text{t},y}, D_{\text{t},z})$) and rotational diffusion constants ($\mathbf{D}_{\text{r}}(D_{\text{r},x}, D_{\text{r},y}, D_{\text{r},z})$) computed from the template radius $R$ (section 1.1). Given $R$, we assume the diffusion constant is isotropic and use the standard Stokes–Einstein relations:

$$\begin{cases} D_{\mathrm{t}} = D_{\mathrm{t},i} = \dfrac{k_{\mathrm{B}}T}{6\pi\eta R} \\ D_{\mathrm{r}} = D_{\mathrm{r},i} = \dfrac{k_{\mathrm{B}}T}{8\pi\eta R^3} \end{cases} \qquad (i \in \{x, y, z\}) \tag{M12}$$

Where $T$ is the temperature, which we use 298.15 K, and $\eta$ is the viscosity of the solution, which we estimate as the viscosity of water at 298.15 K here ($8.9 \times 10^{-4}\ \mathrm{Pa} \cdot \mathrm{s}$).

##### 1.5. Estimate kinetic constants

Using either a default free energy $\Delta G = -16 k_{\mathrm{B}} T$ or the free energy estimated from ProAffinity-GNN, $\Delta G_{GNN}$, the equilibrium constant $K_{\mathrm{eq}}$ is:

$$K_{\mathrm{eq}} = \frac{1}{c_0} e^{-\frac{\Delta G}{k_{\mathrm{B}} T}} \tag{M13}$$

Where $c_0$ is standard concentration defined as:

$$c_0 = 1\ \mathrm{M} = 0.6022\ \mathrm{nm}^{-3} \tag{M14}$$

We assume $T = 298\ K$ in ProAffinity-GNN inference. Consider the Smoluchowski diffusion–limited association-rate:

$$k_{\mathrm{diff}} = 4\pi (D_A + D_B) R_{\mathrm{enc}} \tag{M15}$$

Here let us assume $R = R_A \approx R_B$ and we arbitrarily take 1% of that to assume a value of an on rate that is below the Smoluchowski diffusion limit. Using Stokes-Einstein relations to estimate the diffusion constants, then we have the estimated $k_{\mathrm{a}}$:

$$k_{\mathrm{a}} = 4\pi \left(\frac{k_{\mathrm{B}}T}{3\pi\eta R}\right) \left(\frac{1}{100}(2R)\right) = \frac{4 k_{\mathrm{B}} T}{150\eta} \tag{M16}$$

Note that under this symmetric assumption $R = R_A \approx R_B$, $R$ is eliminated from the estimation, so we basically set $k_\mathrm{a}$ to be constant

$$k_\mathrm{a} = \frac{4k_\mathrm{B}T}{15\eta} \approx 120\mathrm{nm}^3/\mu\mathrm{s} \tag{M17}$$

or $7.4 \times 10^7\ \mathrm{M}^{-1}\mathrm{s}^{-1}$. We vary $k_\mathrm{b}$ as:

$$k_\mathrm{b} = \frac{k_\mathrm{a}}{K_\mathrm{eq}} = \frac{c_0 k_\mathrm{a}}{e^{-\frac{\Delta G}{k_\mathrm{B}T}}} \approx (7.4 \times 10^7\ \mathrm{s}^{-1})e^{\frac{\Delta G}{k_\mathrm{B}T}} \tag{M18}$$

When ProAffinity-GNN failed predicting $\Delta G$ or users configure to skip affinity prediction, we assume a relatively strong $\Delta G = -16k_\mathrm{B}T \approx -40\ \mathrm{kJ/mol}$. In this case, $k_\mathrm{b} \approx 8.2\ \mathrm{s}^{-1}$.

#### 2. Only for repeated subunits: regularize repeated chains

If our complex contains repeated chains, we need to regularize them to assign a single CG geometry and set of interactions, reducing from $N_\mathrm{chain}$ to $N_\mathrm{mol}$ total subunits, and from $N_\mathrm{edge}$ to $N_\mathrm{rxns}$ unique binding interactions. To again use the HIV Gag example (5L93), the $N_\mathrm{edge} = 36$ in the PDB structure are reduced to $N_\mathrm{rxns} = 3$. To do so, we generate a molecular template for each chain group and an interface template for each of its interacting partners. The overall workflow includes (1) building molecule templates, (2) deduplicating interface templates via geometric signatures, (3) instantiating concrete interfaces once per observed contact, (4) aligning all repeated chains back to a common reference frame, and then (5) inferring mutually exclusive interfaces based on steric clashes.

Two local variables are important to keep in mind in this subpackage. First, interface template coordinates are stored as interface template offsets relative to the owning molecule template's COM $\vec{s}$, making templates portable across repeated subunits. Second, interface templates are keyed by geometric signatures (see 1.3a below) normalized to suppress floating-point error to deduplicate templates.

##### 2.1. Iterate repeated groups and build templates

Consider a subunit that has $N_{\text{rep}}$ copies in the assembly. At this point we are working with our rigid body subunits, but for each interface we have retained information about the residues used to form the interface. The first subunit $i = 1$ defines the molecular template. It is possible for repeated subunits to not always have the same number of interfaces as one another from the PDB file, for example if the structure is only a fragment of a complete assembly (like the HIV lattice). More unusually, repeated subunits can have distinct molecular 'neighborhoods' with distinct partners. Most commonly, the subunits have the same sets of interfaces as one another, but not at the exact same positions due to thermal noise.

Given two repeats, template $i$ and another subunit $j \in 2{:}\,N_{\text{rep}}$, we need to identify which interfaces are the same, to determine if one subunit has more interfaces or a distinct set of interfaces. To do so, we use the position of the interface in each subunit, specifically its distance, and we check if its partner is of the same protein type in both cases. Then we verify that the orientation of the contact is the same, at which point we establish they are indeed the same interface. Repeat for all interfaces in each subunit until we have established a minimal unique set of interfaces. For all chains $j$, we overwrite their internal interface lists with the templates list, so that they can be aligned properly in subsequent steps. We repeat this for all chains $j$ in the repeated group. The template is updated if new interfaces are found.

For a contact between subunit $i$ and its partner $j$, define COMs and interface coordinates:

$$\vec{c}_i, \vec{c}_j, \vec{s}_a^i, \vec{s}_b^j \in \mathbb{R}^3 \tag{M7}$$

Geometric signatures are summarized by the tuple $\mathbf{geo}(i,j)$:

$$\mathbf{geo}(i,j) = \left(d_i, d_j, \theta_i, \theta_j\right) \tag{M8}$$

Where $d_i, d_j$ are the COM-to-interface distances and $\theta_i, \theta_j$ are the angles between COM-to-interface vector and the COM-to-COM vector, i.e. let $\vec{s}_a^i$ be the offset vector defined as $\vec{s}_a^i = \vec{s}_a - \vec{c}_i$, then the signatures $\mathbf{geo}(i,j) = \left(d_i, d_j, \theta_i, \theta_j\right)$ are defined as:

$$\begin{cases} d_i = \left\|\vec{s}_a^i\right\| \\ d_j = \left\|\vec{s}_b^j\right\| \\ \theta_i = \frac{\vec{s}_a^i \cdot (\vec{c}_j - \vec{c}_i)}{\|\vec{s}_a^i\| \|\vec{c}_j - \vec{c}_i\|} \\ \theta_j = \frac{\vec{s}_b^j \cdot (\vec{c}_i - \vec{c}_j)}{\left\|\vec{s}_b^j\right\| \|\vec{c}_i - \vec{c}_j\|} \end{cases} \tag{M9}$$

Note that because we assume that there is at most one interaction per pair of subunits, any $(i,j)$ pair maps uniquely to an $(a,b)$ pair, which is the reason why $\mathbf{geo}$ only requires input of chain pair $(i,j)$.

If subunit $i, j$ are the same type, then they are homodimers. In this case, we need to proceed with special caution: we need to distinguish between homodimeric heterotypic (asymmetric) interactions from homodimeric homotypic (symmetric) interactions.

##### *2.1a. Homodimeric homotypic interactions*

We classify the interaction as homodimeric homotypic if the signature is nearly symmetric:

$$|d_i - d_j| < \delta_d \ \ \mathbf{and}\ \ |\theta_i - \theta_j| < \delta_\theta \qquad \text{(M10)}$$

With $\delta_d$ the internal distance threshold and $\delta_\theta$ the angle threshold (both are specified in hyperparameters). A homotypic interaction uses one interface template shared by both sides, i.e. both site $a$ and $b$ are of the same site template. They will eventually be regularized to using the identical geometry.

##### *2.1b. Homodimeric heterotypic interactions*

If the threshold check fails above, we create two interface templates respectively for $a$ and $b$, each stored with its corresponding signatures: signature for $a$ is $\mathbf{geo}(i,j) = (d_i, d_j, \theta_i, \theta_j)$ and signature for $b$ is $\mathbf{geo}(j,i) = (d_j, d_i, \theta_j, \theta_i)$.

##### *2.1c. Determine existing interface templates*

When determining the interface template of a candidate pair of subunits, we first determine whether any existing template already has the same pair of interaction subunit types as the candidate. If not, then the candidate pair of subunits is assigned to a new chain type. If some existing template has the same pair of interaction subunit types, the we test the similarity between the geometric signatures: we compare the geometric signature of the template $\mathbf{geo}_{\mathrm{temp}}(k,l) = (d_k, d_l, \theta_k, \theta_l) =$ against the candidate $\mathbf{geo}_{\mathrm{cand}}(i,j) = (d_i, d_j, \theta_i, \theta_j)$, where $\delta_d$ is the internal distance threshold and $\delta_\theta$ is the angle threshold:

$$|d_k - d_i| < \delta_d \ \ \mathbf{and} \ \ |d_l - d_j| < \delta_d \ \mathbf{and} \ \ |\theta_k - \theta_i| < \delta_\theta \ \mathbf{and} \ \ |\theta_l - \theta_j| < \delta_\theta \qquad \text{(M11)}$$

If the similarity test (equation M11) passes, we assign the candidate interaction as the same type as the existing template. Otherwise, we create a new interaction template for the candidate. We then build binding interface templates for each molecule templates based on the reference chain of the group (its index within the group is 0). Before adding, we check if an interface with the given name exists in the molecule's interface list to avoid creating duplicates when the same edge is visited again from the other direction or another pass.

There are special cases as noted in the main text where a repeated subunit has different partners in different parts of the complex. Here, we pause and check whether the reference chain contains all the interface types of this molecule type. This is to accommodate cases where molecules of the same type are in different binding states. We proceed if the reference chain contains all the interface types. Otherwise, we come up with a template that contains all possible interfaces. This is done by iterating through all molecule instances of the same type with binding sites not existing in the reference chain, calculating a Kabsch transformation via the interfaces that exist in both the reference chain and the iterated chain, and using that Kabsch transformation to project the iterated chain onto the reference chain. From the result, we can add the interfaces that are initially present to the reference chain. The new template containing all the interfaces is called "mixed-frame instance" in ioNERDSS. Note that this might be unintentional when the interfaces are realistically the same type but slightly misaligned in the PDB. If $\delta_d$ and $\delta_\theta$ are too strict ioNERDSS recognizes violations of the thresholds as different chains. Therefore, whenever a mixed-frame instance is created, ioNERDSS sends out a warning to the user to make sure that it is intended (usually benign). For example:

```
ionerdss.pdb.XXXX - WARNING - Representative instance 6534397760 missing interfaces:
['A1', 'L1', 'K2']. Falling back to mixed-frame instances (RISKY).
```

##### 2.2. Propagate template geometry across repeats

After all templates and concrete interfaces are in place, we push the reference template geometry into every member of each repeated group. For the group's reference chain we keep its COM and define a body-fixed *reference coordinate* that is assigned to each subunit *per interaction,* but for simplicity is typically set to $\vec{s}^{\,i}_{ref} = [0,0,1]$, as its only restriction is that it is not co-linear with an interface vector. For each non-reference member, we compute the Kabsch transformation from the reference chain to the member and apply it to the reference COM and to each template offset converted back to absolute coordinates. The transformed COM and interface coordinates are written back into the corresponding chains and interfaces. This guarantees that within a group all members share a coherent geometry consistent with the reference. An updated CG model with the regularized chain COMs and interface coordinates is created by deep-copying the original CG model and replacing the regularized chain COMs and interface coordinates with the regularized values taken from the molecules, preserving the original partner order per chain; if a partner is missing (error handling), we fall back to the original coordinate.

##### 2.3 Optional: Check Steric clashes to infer mutually exclusive interfaces

An optional case is to test if two interfaces cannot be both occupied at the same time, despite that they both represent two unique interfaces, with an example being the GATOR complex. From a modeling perspective, we want a reaction network that excludes the second interface from binding when the first is already bound, to enforce this steric effect. This can be written as conditional binding reactions in the reaction file, such that if one site is occupied, the other is forbidden from reacting. We infer such exclusions without simulation by explicit Cα-level clash checks. For a candidate interface on chain $C_i$ binding to $C_j$ we extract the Cα coordinates of $C_j$ and transform them into the frame of a previously processed chain $C_i{}'$ within the same group (i.e. $C_i$ and $C_i{}'$ are repeated chains) using the Kabsch rigid transform between $C_i$ and $C_i{}'$. We then iterate through the set of all other registered binding partners of $C_i{}'$ to test the transformed set against the Cα coordinates. If a Cα clash is detected, we mark the interface template as mutually exclusive by adding each template's full name to the other's "required free list", denoting the steric incompatibility. To avoid quadratic blow-ups, we only perform this test on the first appearance of a given molecule template within the group.

#### 3. Predicting binding affinity using ProAffinity-GNN

ProAffinity-GNN[1] estimates interface binding affinities by combining structure aware graph attention neural networks with protein language model embeddings. It achieves high correlation and low deviation (R=0.669, MAE=1.50 kcal/mol). For each protein–protein complex, residues at the contact interface are represented as nodes with geometric, physicochemical, and learned sequence features, while edges encode inter residue proximity/orientation. Iterative message passing captures multi body coupling and context across the interface. Global pooling layers then aggregate residue level signals to a complex level affinity, allowing the model to leverage both local patch chemistry and longer-range architectural constraints. Trained on a curated dataset of experimentally characterized complexes, the framework generalizes across diverse topologies and oligomerization states and delivers competitive accuracy for structure based binding affinity prediction.

To accept PDB files as inputs, ioNERDSS supports a naïve atom type encoder (Carbon: 'C', oxygen: 'OA', Nitrogen: 'N', Sulfur: 'SA', Hydrogen: 'H', others: 'A'). We also integrate AutoDockFR (ADFR) for labelling atoms and convert PDB files to PDBQT files. Before conversion, we sanitize the PDB files by deleting everything other than conventional amino acids (20 amino acids) and nucleotides (DA, DG, DC, DT and A, G, C, U, T). Using PDBQT files converted by ADFR "prepare_recepter", we achieved good inference accuracy for weak bindings (Fig. SI). The binding affinity are under-estimated for strong binders with the binding affinity $\Delta G < -50\ \mathrm{kJ/mol}$, or $K_{\mathrm{eq}} > 5.76 \times 10^{8}\ \ \mathrm{M}^{-1}$ (Figure S2).

We assume that on-rates are close to but not quite the diffusion limit, we then vary the off-rate such that the equilibrium constant conforms to the free energy predicted by the ProAffinity-GNN (see section SI 1.3d).

#### 4. Determining NERDSS and ODE hyperparameters

While most of the hyperparameters are provided by the user via optional arguments, some of the NERDSS and ODE hyperparameters are automatically determined from other geometric or system parameters (overridable).

##### 4.1. NERDSS time step and ODE time span

In NERDSS software, to limit interactions to two-body association within a time step, the time step is constrained by bimolecular binding radii, reactant densities, and diffusion. Using the FPR[2] (Free-Propagator Reweighting) definition of the maximum diffusive displacement leading to collision, the appropriate time step for simulating with NERDSS binding of B to A $\Delta t_{A,B}$ is given by:

$$\Delta t_{A,B} = \frac{1}{56(D_A + D_B)}\left(\left(\frac{3}{4\pi\rho_A} + {\sigma_{AB}}^3\right)^{\frac{1}{3}} - \sigma_{AB}\right)^2 \qquad \text{(M19)}$$

Where $D_A$ and $D_B$ are translational diffusion constants of correspond species (see section SI 1.3d for definition), $\rho_A$ is total number of A divided by system volume, and $\sigma_{AB}$ is bond length between A and B. The ioNERDSS package iterates through all binding reactions, calculates the limiting time step for each interacting pair (checking both directions A+B and B+A for density dependence) based on the method above, and selects the minimum robust time step for the whole system ($\Delta t_{\text{sys}}$):

$$\Delta t_{\text{sys}} = \min\left(\Delta t_{A,B}\right) \qquad \text{(M20)}$$

The ODE time span is default to align with the time span of NERDSS simulation, which is:

$$(0, N_{\mathrm{itr}} \cdot \Delta t_{\mathrm{sys}}) \tag{M21}$$

Where $N_{\mathrm{itr}}$ is the total number of iterations. This time span can be overridden to a custom fixed value via changing the `ode_time_span` hyperparameter when building the model.

##### 4.2. NERDSS initial count and ODE initial concentration

The NERDSS initial counts of species are set to the total number of molecules partitioned by the species stoichiometry found in the given PDB file. Specifically, a target total molecule count $N_{\mathrm{tot}}$ (defined by `nerdss_total_molecule_count`, default to 75) is distributed among these unique species proportional to their stoichiometry. For a species $i$ with stoichiometry $S_i$, the count $N_i$ is calculated as:

$$N_i = \left\lceil \frac{S_i}{\sum_j S_j} \cdot N_{\mathrm{tot}} \right\rceil \tag{M22}$$

Where ⌈⌉ denotes rounding up. The ODE initial concentrations $C_{\mathrm{init},i}$ are then automatically derived from these calculated particle counts $N_i$ and the simulation box volume $V_{\mathrm{box}}$, ensuring that both the NERDSS simulations and ODE start with physically identical concentrations with the equation below ($N_A$ denotes Avogradro's number):

$$C_{\mathrm{init},i} = \frac{N_i}{N_A V_{\mathrm{box}}} \tag{M23}$$

##### 4.3. NERDSS steric overlap check (overlapSepLimit)

NERDSS uses a simulation parameter termed overlapSepLimit that by default is set to 0.1 nm, to evaluate if new subunits that associate onto an existing complex are sterically clashing with other subunits in the complex. The distance $d_{CC}$ is measured between the new subunit $i$ COM and the COM of all other subunits in the complex. If $d_{CC} < overlapSepLimit$ , the association event is rejected. For heteromeric and defect-free complexes, the distances between all COM pairs is either the same as in the designed structure, $d_{CC} = d_{CC,target}$, or if the new subunit attaches to a free site where another subunit already occupies that space (e.g. if a trimer is 'open' and not yet 'closed), this produces $d_{CC}$ ~0 nm, which is thus correctly rejected due to steric overlap. For structures with repeated subunits that can produce defects or imperfect alignment, we can increase this limit to ensure that two subunits that have $d_{CC} > 0.1$nm but are occupying the same spot are still rejected. We also compute an upper bound for the $overlapSepLimit$ based on the measured COM-COM distances between all pairwise proteins in the target complex, $d_{stericBound} = \min(d_{CC,target})$. This ensures that an associating subunit that binds to an open and sterically available position is not rejected due to users setting $overlapSepLimit$ too high. Our tutorials force this parameter to not exceed this value, $overlapSepLimit \leq d_{stericBound}$.

#### 5. Graph-based ODE system generation

##### 5.1. Convert coarse-grained structure to a simple typed graph

To systematically generate an ODE system, the coarse-grained structural representation is converted into a simple typed graph. Specifically, each molecular subunit or domain is represented as a node, and physical interaction (mediated by interfaces) between subunits are represented as edges labeled by interaction type. Symmetric assemblies with repeated subunits are reduced to symmetric graphs with repeated node and edge types to preserve the topology (see Figure M1).

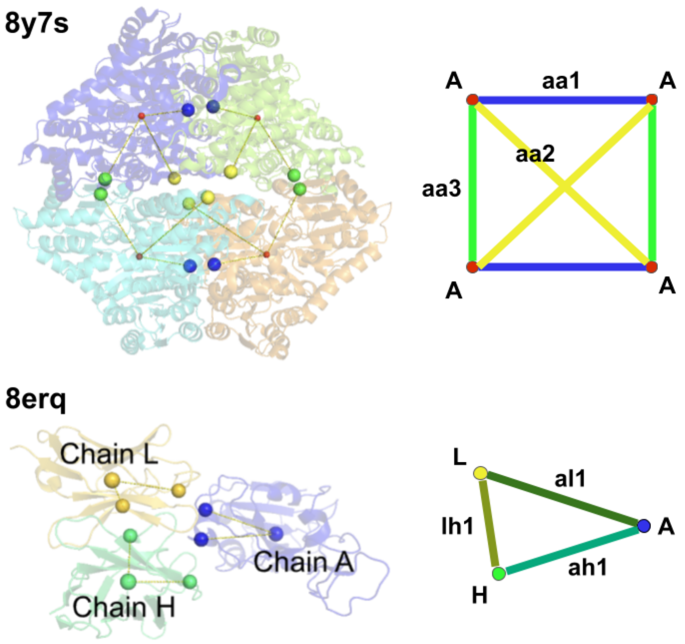

**Figure M1.** Coarse-grained (left) and graph (right) representations of two example protein complexes showing subunits and interaction sites. Top: a homomeric assembly (PDB ID 8y7s) represented by identical node types and multiple interface types (aa1–aa3). Bottom: a heteromeric complex (PDB ID 8erq) represented by distinct node types (A, H, L) and interaction types (al1, ah1, lh1). In general, each new edge per node indicates a new interface.

##### 5.2. Induction based subgraph

Based on the graph that represents the full assembly, we generate all possible intermediates via graph induction, which is defined as the subgraph obtained by removing one or more nodes (in sequence) from the original graph together with all edges connected to those nodes. Each induced subgraph therefore corresponds to a partial assembly composed of a subset of the original subunits while preserving the interaction among the remaining components (Figure M2).

Systematically enumerating induced subgraphs provides a complete set of candidate assembly intermediates that are topologically consistent with the full complex. To avoid double counting the intermediates in the symmetric pathways (e.g. with repeated subunits), we consider all nonempty subsets of nodes and construct the corresponding induced subgraphs, followed by deduplication under graph isomorphism while respecting node and edge types. This removes redundancies arising from symmetry or relabeling and yields a list of unique intermediates. Furthermore, the reactions along the assembly pathway can be viewed as a sequence of node additions or removals that transform one induced subgraph into another.

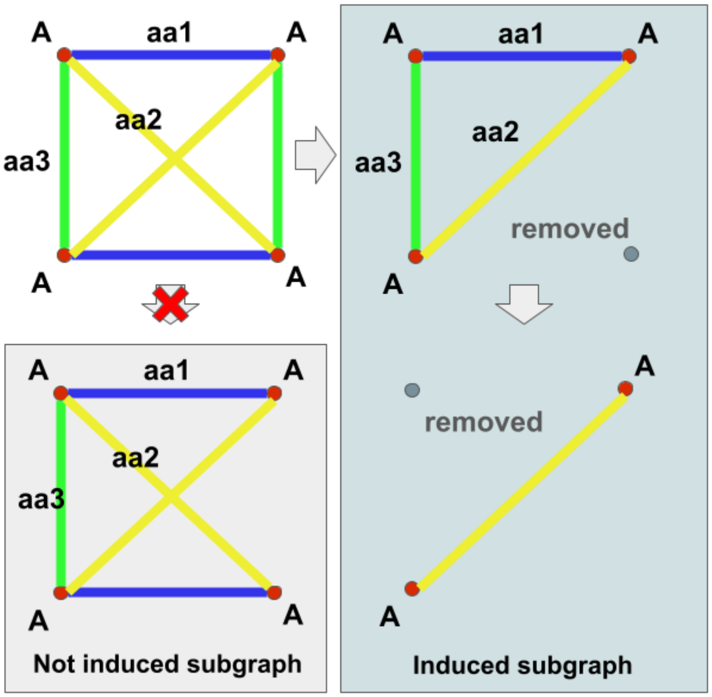

**Figure M2 Induced subgraph versus not induced subgraphs.** Right: sequential removal of nodes produces valid induced subgraphs that preserve all interactions among the remaining nodes. Left bottom: example of a graph that is not an induced subgraph, because some edges between retained nodes are omitted. Induced subgraphs therefore maintain the full connectivity structure among the surviving nodes while representing physically consistent assembly intermediates.

Species represented by subgraphs and reactions obtained via this method ensures that the reaction network contains only second order reactions, where ring closing such as in Figure M2 is concatenated with binding of the last subunit into one effective reaction. We note that NERDSS simulations do not operate via dissociation or association events that break/form all edges per subunit. Each association or dissociation event in NERDSS involves only a single edge. As we discuss in the main text, because co-localized subunits then form remaining bonds using first-order reactions, the free energies are the same, and the kinetics can also be designed to agree with high accuracy.

##### 5.3. Mathematical formulation of intermediate enumeration problem

Let $\mathcal{T}_V$ be a finite set of vertex types (representing subunit types) and $\mathcal{T}_E$ be a finite set of edge types (representing interaction types). Define a graph $\mathcal{G} = (V, E, \tau_V, \tau_E)$ where:

- $V$ is a finite set of vertices.
- $E \subseteq \{\{u, v\} | u, v \in V, u \neq v\}$ is a set of undirected edges (ensuring no loops or multi-edges).
- $\tau_V: V \to \mathcal{T}_V$ assigns one vertex type to each vertex and
- $\tau_E: E \to \mathcal{T}_E$ assigns one edge type to each edge.

The graph equivalence can be defined as: two graphs $\mathcal{G}_1 = \left(V_1, E_1, \tau_{V_1}, \tau_{E_1}\right)$ and $\mathcal{G}_2 = \left(V_2, E_2, \tau_{V\,2}, \tau_{E\,2}\right)$ are equivalent if there exists a bijection

$$\phi: V_1 \to V_2$$

Such that

- $\tau_{V_1}(v) = \tau_{V_2}\left(\phi(v)\right)$ for all $v \in V_1$
- $\{u, v\} \in E_1 \Leftrightarrow \{\phi(u), \phi(v)\} \in E_2$
- $\tau_{E_1}(\{u, v\}) = \tau_{E_2}(\{\phi(u), \phi(v)\})$ for all $\{u, v\} \in E_1$

Which ensures both structure and typing are preserved in an equivalence relation.

Similarly, we can define induced subgraph: a graph $\mathcal{H} = \left(V_{\mathcal{H}}, E_{\mathcal{H}}, \tau_{V_{\mathcal{H}}}, \tau_{E_{\mathcal{H}}}\right)$ is an induced subgraph of $\mathcal{G} = (V, E, \tau_V, \tau_E)$ (noted $\mathcal{H} \subseteq \mathcal{G}$) if:

- $V_{\mathcal{H}} \subseteq V$
- $E_{\mathcal{H}} \subseteq \left\{\{u, v\} \in E \middle| u \in V_{\mathcal{H}}, v \in V_{\mathcal{H}}\right\}$
- $\tau_{V_{\mathcal{H}}} = \tau_V | V_{\mathcal{H}}$, i.e. the restriction of $\tau_V$ to $V_{\mathcal{H}}$
- $\tau_{E_{\mathcal{H}}} = \tau_E | V_{\mathcal{H}}$

Trivially, $\mathcal{G} \subseteq \mathcal{G}$ and $(\emptyset, \emptyset, \emptyset \to \emptyset, \emptyset \to \emptyset) \subseteq \mathcal{G}$. Now the question is how to efficiently get all unique non-trivial induced subgraphs of $\mathcal{G}$.

##### 5.4. Implementation of Avis-Fukuda reverse searching to get all unique non-trivial induced subgraphs

To efficiently enumerate the unique, non-trivial connected induced subgraphs of $\mathcal{G}$, we combine Avis-Fukuda reverse search[3] with Weisfeiler-Lehman (WL) graph hashing, inspired by previous research on optimized enumeration on self-assembly pathways[4]. Rather than generating the full powerset of node subsets and then filtering for connectivity, the algorithm traverses only the space of connected induced subgraphs. For each connected component of $\mathcal{G}$, a reverse-search tree is defined in which each connected induced subgraph $S$ has a unique parent $P(S)$, obtained by removing the largest-index removable vertex while preserving connectivity. Starting from singleton vertices, the search expands a subgraph only by adding neighboring vertices $y$ such that $P(S \cup \{y\}) = S$, which guarantees that every connected induced subgraph is visited exactly once without explicit powerset enumeration. For each visited subgraph, we compute a 1-dimensional WL hash using both node attributes (subunit identities) and edge attributes (interface bond types). This hash serves as an isomorphism-aware signature, allowing topologically equivalent subgraphs arising from repeated or symmetric subunits to be deduplicated efficiently. Subgraphs with previously unseen WL hashes are retained as unique species, while isolated singleton nodes are excluded as trivial cases. In this way, the method yields the set of unique assembly intermediates while substantially reducing the combinatorial cost of subgraph enumeration.

##### 5.5. Reaction enumeration using the set of intermediate species

Having established the complete, deduplicated set of intermediate species $S = \{\mathcal{G}_S | \mathcal{G}_S \subsetneq \mathcal{G}, \mathcal{G} \neq (\emptyset, \emptyset, \emptyset \rightarrow \emptyset, \emptyset \rightarrow \emptyset)\}$ via Weisfeiler-Lehman hashing, the next goal is to connect these states through a network of reactions. Because our graph building method guarantees that any assembly event can be broken down into individual binding steps (or concatenated effective steps), we restrict the network to second-order association and first-order dissociation reactions.

To systematically construct the list of reactions, we evaluate every candidate product species $\mathcal{G}_S \in S$. For each $\mathcal{G}_S$, we iterate through all possible ways to partition its nodes into two non-empty, disjoint, and fully connected subgraphs $\mathcal{G}_A$ and $\mathcal{G}_B$. Practically, it is done via iterating through $\mathcal{G}_A \subseteq \mathcal{G}_S$ and check if $\mathcal{G}_B = \mathcal{G}_S - \mathcal{G}_A$ is fully connected and non-trivial by checking whether their WL hash existed in our set of WL hashes. If both $\mathcal{G}_A$ and $\mathcal{G}_B$ exist in our unique species set $S$ identified via their invariant WL hashes, we define a reversible bimolecular reaction: $\mathcal{G}_A + \mathcal{G}_B \rightleftharpoons \mathcal{G}_S$. This process guarantees that we capture all valid assembly pathways up to the given full assembly, while some reactions are included using one effective reaction representing the concurrent formation of multiple interfaces, such as the ring closing reaction.

##### 5.6. Estimate the reaction rate constants of the effective reactions

To simulate the assembly kinetics across this enumerated network, we must estimate the macroscopic rate constants for each effective reaction that combines several fundamental steps (such as the ring closing step described above in section 4b). The forward association rate for a simple bimolecular binding event is approximated to be a constant (see section 1.3d) using diffusion-limited collision theory, scaled by a statistical degeneracy factor that accounts for the number of equivalent binding orientations (here analogously evaluated by counting graph isomorphisms) between the reactants and the product subgraph. The corresponding reverse dissociation rate is then calculated via detailed balance, using the binding free energy of the specific interface. Specifically, to assign the corresponding dissociation rate $k_{\text{off}}$, we enforce thermodynamic detailed balance. Let $G(\mathcal{G})$ denote the total Gibbs free energy of the internal bonds within a subgraph $\mathcal{G}$, computed as the sum of the interaction energies $G(\mathcal{G}) = \sum G_E$ of all its edges. The standard free energy of the reaction is the energy associated with the novel interfaces formed when $\mathcal{G}_A$ and $\mathcal{G}_B$ bind to form $\mathcal{G}_k$: $\Delta G_{\text{rxn}} = G(\mathcal{G}_k) - \big(G(\mathcal{G}_A) + G(\mathcal{G}_B)\big)$. The macroscopic equilibrium constant $K_{\text{eq}}$ is determined by: $K_{\text{eq}} = e^{-\frac{\Delta G}{RT}}$. The dissociation rate is computed directly from the forward rate and the equilibrium constant. This explicit calculation also handles effective structural ring closures. When the binding of $\mathcal{G}_A$ and $\mathcal{G}_B$

closes a ring, $\Delta G_{\mathrm{rxn}}$ sums the free energy of all bonds formed and thus captures the total energy of all simultaneously formed interfaces. While the forward binding rate $k_{\mathrm{on}}$ remains fixed and determined by diffusion (1.3d), the resulting $k_{\mathrm{off}}$ is calculated from the free energy and the binding rate. Importantly, with this method, for effective reactions that involve a binding event concatenated with rapid structural ring closing, the forward rate remains largely governed by the initial diffusional encounter, assuming the second ring closing step is much faster. However, the effective reverse rate is substantially reduced, reflecting that breaking the ring requires two consecutive dissociation step which becomes much harder.

#### 6. Example use cases of ioNERDSS

The raw data of all the examples in the main text are collected in the following GitHub repository:

https://github.com/JohnsonBiophysicsLab/ionerdssPaperSI

##### 6.1. Ideal dodecahedron assembly simulation

The simulation is initiated with the `ioNERDSS.PlatonicSolids` module. The radius of the dodecahedron is 5 nm.

```
Kinetic constants
D = [13.0, 13.0, 13.0]
Dr = [0.3, 0.3, 0.3]
onRate3Dka = 10
offRatekb = 10000
```

```
Parameters
timestep = 0.1
monomerCount = 1300
WaterBox = [350.0, 350.0, 350.0] # nm
```

The NERDSS simulation results is available at

github.com/JohnsonBiophysicsLab/ionerdssPaper/tree/main/dodecahedral_dir/nerdss_files

##### 6.2. actin filament (6BNO[5]) assembly simulation

This example is available on GitHub at

https://github.com/JohnsonBiophysicsLab/ionerdss/blob/main/tutorials/ionerdss_tutorial_6bno.ipynb

with the following set of parameters:

```
Kinetic constants
onRate3Dka = 10
#offRatekb: predicted by proaffinity-gnn
```

```
Parameters
timeStep = 0.4
monomerCount = 130
WaterBox = [600, 600, 600] # nm
```

To use the same set of parameters as above, replace the `build_model_from_pdb` step with the following code (changing the optional arguments; a full list of available parameters is provided in Table SA). Be aware that this requires ADFR installed and proaffinity-gnn configured on the local machine.

```
system = build_system_from_pdb(
    source=pdb_id,
    workspace_path=f"{pdb_id}_dir",
    # Interface detection
    interface_detect_distance_cutoff=1.5,
    interface_detect_n_residue_cutoff=5,
    chain_grouping_seq_threshold=0.5,
    nerdss_water_box=[600.0, 600.0, 600.0],
    default_on_rate_3d_ka=10,
    nerdss_time_step=0.4,
    nerdss_totoal_molecule_count=130,

    # ProAffinity-GNN
    predict_affinity=True,            # Enable proaffinity-gnn
    adfr_path=”REPLACE_WITH_PATH_TO_ADFR”,      # Path for ADFR software

    # ODE Pipeline Configuration
    ode_enabled=True,             # Now using System-compatible generator!
    ode_time_span=None,    # Auto-calculated based on NERDSS simulation time
```

```
    ode_solver_method="BDF",     # Solver for stiff systems
    ode_plot=True,               # Generate plots
    ode_save_csv=True,            # Save data to CSV

    # Transition matrix parameters
    count_transition=True,        # Enable transition matrix tracking
    transition_matrix_size=100,   # Size of matrix (max cluster size expected)
    transition_write=1000,         # Write every 1000 iterations
)
```

##### 6.3. HIV-1 CA-SP16 (5l93) assembly simulation

This example is available on GitHub at
https://github.com/JohnsonBiophysicsLab/ionerdss/blob/main/tutorials/ionerdss_tutorial_5l93.ipynb

##### 6.4. Benzaldehyde lyase mutant M6 from Herbiconiux sp. SALV-R17 (8y7s) assembly simulation

This example is available on GitHub at

https://github.com/JohnsonBiophysicsLab/ionerdss/blob/main/tutorials/ionerdss_tutorial_8y7s.ipynb

### SUPPORTING FIGURES

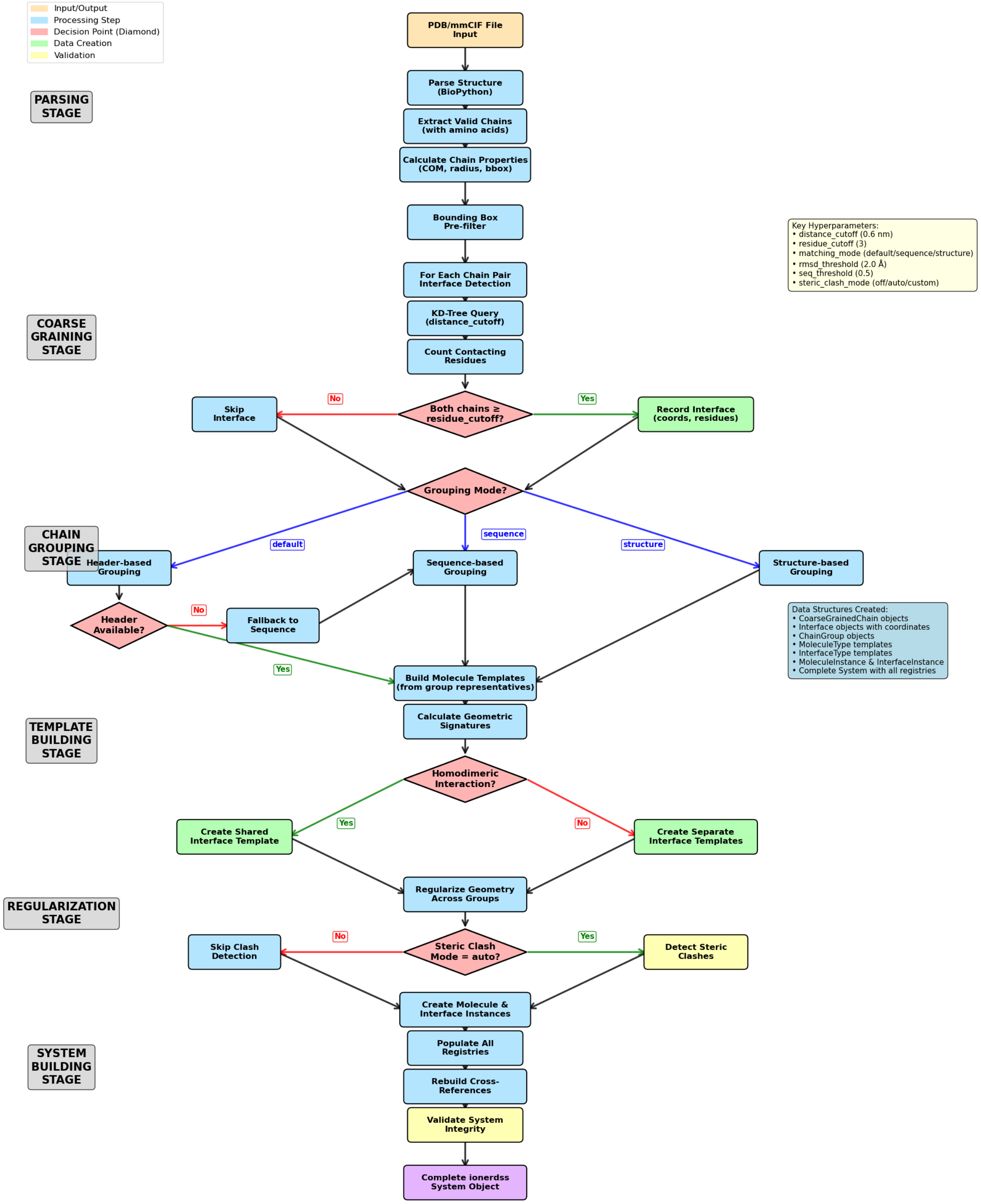

**Figure S1. Flowchart for PDB to NERDSS parameter pipeline.**

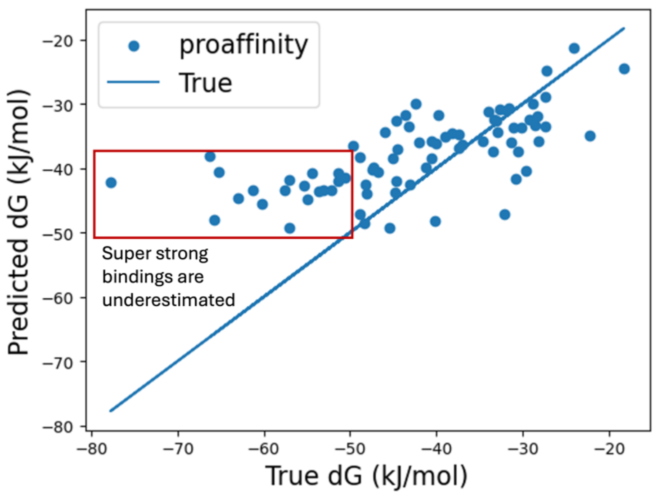

**Figure S2. Predictive performance of easy-to-use ProAffinity-GNN inference.** This lightweight implementation is not as accurate as the full package. Specifically, sequences are not aligned with canonical FASTA sequences. However, the binding affinity inference still agrees reasonably well with the true binding free energies of the test set used in the original study.

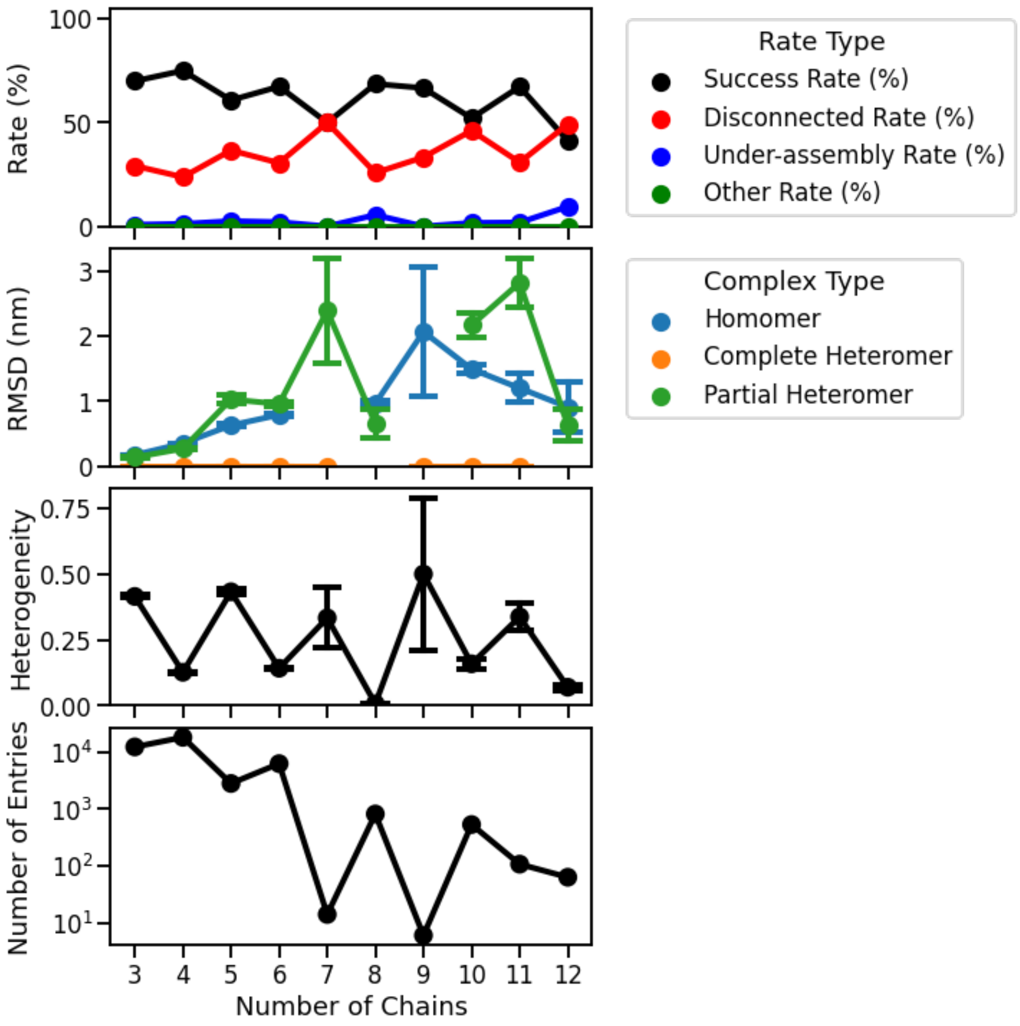

**Figure S3. Success rate and average RMSD of validating ioNERDSS PDB coarse-graining to reaction-diffusion simulation pipeline and average heterogeneity and number of the input PDB entries.** 44703 PDBs were selected and tested based on criteria of protein-only entries, experimental methods (X-ray diffraction or electron microscopy), exact protein chain count (protein entity count and number of chains matches in the PDB entry), resolution (<= 3.5 Å), size (number of chains between 3 and 12). Validation simulations are run with default hyperparameters, except for a smaller box size (50 nm x 50 nm x 50 nm) and a lower initial copy numbers (enough to assemble 1 copy of full structure encoded in the PDB). RMSD is calculated only for the succeeded cases between the COM of design model and the NERDSS assembled structure. Success rate decreases overall with larger assemblies with high fluctuations and RMSD increases with larger assemblies up to 10 but decreases after that. Heterogeneity is defined as (number of chain types – 1) / (number of chains – 1), which is 0 for homomers and 1 for full heteromers. Note that heterogeneity has a clear even-odd switching pattern which may contribute to the fluctuation of success rates. Error bars represent ±1 standard error.

#### SUPPORTING TABLES

| Parameter | Default | Description |
|---|---|---|
| **1. Core interface detection** | | |
| `interface_detect_distance_cutoff` | 0.6 nm | $r_{\mathrm{cut}}$: Max distance between atoms to consider as interface |
| `interface_detect_n_residue_cutoff` | 3 | $N_{\mathrm{cut}}$: Min contacting residues per chain to validate interface |
| `min_chain_length` | 4 residues | Min residues for a chain to be included |
| **2. Core interface detection** | | |
| `homotypic_detection` | `"auto"` | Mode: `"auto"`, `"signature"`, or `"off"` |
| `homotypic_detection_interface_radius` | 8.0 Å | Detection radius for homotypic binding |
| `3` | 0.5 | Biopython similarity score for homotypic binding |
| `homodimer_distance_threshold` | 0.5 nm | Distance threshold for homodimer detection |
| **3. Chain grouping & alignment** | | |
| `chain_grouping_seq_threshold` | 0.5 | $\delta_{\mathrm{seq}}$: Sequence identity threshold for grouping |
| `chain_grouping_matching_mode` | `"default"` | `"default"`, `"sequence"`, or `"structure"` |
| `chain_grouping_custom_aligner` | None | Custom `Bio.Align.PairwiseAligner` object |
| **4. Geometry & regularization** | | |
| `is_on_sphere` | `"False"` | Enable spherical alignment when set to`"True"` |
| `steric_clash_mode` | `"off"` | `"off"`, `"auto"` |
| `signature_precision` | 6 | Decimal precision for geometric signatures |
| `template_regularization_strength` | 0.0 | Strength for template fitting regularization |
| `interface_type_assignment_distance_threshold` | 2.0 Å | $\delta_d$:Distance threshold for merging interface types |
| `interface_type_assignment_angle_threshold` | 0.2 rad | $\delta_\theta$: Angle threshold for merging interface types |
| **5. NERDSS simulation setting (*)** | | |
| `generate_nerdss_files` | `True` | Generate simulation files if set to true |
| `nerdss_total_molecule_count` | 75 | Total target molecules (stoichiometry-based; SI 3b) |
| `nerdss_water_box` | `[500, 500, 500]` | Box dimensions in nm |
| `nerdss_time_step` | None | Simulation timestep (us). Auto-calculate (SI 3a) if None. |
| `nerdss_n_itr` | 100,000 | Number of simulation steps |
| `default_on_rate_3d_ka` | 120.0 | Default microscopic binding rate ($nm^3$/us) (SI 1.3d) |
| **6. ODE pipeline parameters** | | |

| | | |
|---|---|---|
| `ode_enabled` | `False` | Enable ODE calculation |
| `max_complex_size_ode` | 12 | Max complex size for ODE generation |
| `ode_time_span` | None | Time span (s). Auto-calculate from NERDSS params if None (SI 3a). |
| `ode_solver_method` | `"BDF"` | SciPy Solver used in solving reaction ODE: `"BDF"`, `"LSODA"`, etc. |
| `ode_atol` | 1e-4 | Absolute tolerance for ODE solver |
| `ode_plot` | `True` | Generate plots when set to true |
| `ode_save_csv` | `True` | Save ODE solution as CSV data |
| `ode_plot_species_indices` | None | Specific species indices to plot; plot all species when set to None (default) |
| `ode_plot_sample_points` | 1000 | Number of time points for plots and csv |
| `ode_species_labels` | None | Custom labels for plots |
| **7. NERDSS simulation: transition matrix parameters** (**WARNING****) | | |
| `count_transition` | `False` | Enable transition matrix tracking |
| `transition_matrix_size` | 500 | Max cluster size to track in matrix |
| `transition_write` | None | Write interval (defaults to nItr/10) |
| **8. ProAffinity-GNN Parameters** (**WARNING*****) | | |
| `predict_affinity` | `False` | Enable ProAffinity-GNN binding energy prediction |
| `adfr_path` | None | Path to ADFR `prepare_receptor` tool |
| **9. Other parameters** | | |
| `generate_visualizations` | `True` | Generate visualization outputs |
| `pdb_file_format` | `"bioassembly 1"` | File / format to fetch from the PDB website |

**Table S1. Physical parameters and simulation hyperparameters used in the ioNERDSS pipeline.** (*) Parameter names conform to NERDSS parameter files.[8] (**) Requires tracking to be enabled (`count_transition=True`). A safe `transition_matrix_size` MUST be specified (recommended to be = `nerdss_total_molecule_count`), otherwise the simulation will error out if the transition matrix is not big enough to hold the largest cluster). (***) Requires installing the ADFR package and specifying the `adfr_path` below. If not using ProAffinity-GNN to predict energy, the code automatically sets the binding free energy for each pair of interfaces to $-16RT$. Users can manually change these values in the output `parms.inp`.

| Metric | Overall | Homomer | Complete Heteromer | Partial Heteromer |
|---|---|---|---|---|
| %***Success*** | 70.09% | 76.56% | 58.82% | 61.92% |
| %***FP*** | 2.46% | 3.98% | 0.33% | 0.00% |
| %***DC*** | 26.15% | 17.80% | 40.07% | 37.35% |
| %***NC*** | 0.00% | 0.00% | 0.00% | 0.00% |
| %***UA*** | 1.30% | 1.66% | 0.79% | 0.73% |
| %***OA*** | 0.00% | 0.00% | 0.00% | 0.00% |
| Avg. RMSD in Success | 0.3327 nm | 0.3814 nm | 0.0000 nm | 0.4828 nm |
| Stdev. RMSD in Success | 0.6728 nm | 0.6554 nm | 0.0000 nm | 0.9264 nm |
| < 1 nm RMSD % out of Success | 85.47% | 82.55% | 100.00% | 81.97% |
| < 0.1 nm RMSD % out of Success | 67.95% | 61.09% | 100.00% | 61.86% |

**Table S2. Validation result for ioNERDSS PDB coarse-graining and simulation pipeline. *Success*** indicates that the target assembly (composition only, check RMSD for geometry) was found and RMSD of COM was computed successfully. ***FP*** (few proteins): Too few protein chains (0 or 1) remain after coarse graining. We ignore anything that has less than 4 amino acids (considered as non-protein species or small molecules). These are guaranteed to fail assembly. ***DC*** (disconnected): The designed assembly graph is disconnected, so the system cannot form the intended single target N-mer. Changing the hyperparameters helps in this case. ***NC*** (NERDSS crashed): NERDSS crashed during export or simulation, such as a segfault, abort, core dump, or similar runtime failure. ***UA*** (under-assembly): The target assembly was not found, and the largest observed assembly is smaller than the target assembly size. ***OA*** (over-assembly): The target assembly was not found, and the largest observed assembly is at least as large as the target assembly size. 44703 PDBs were selected and tested. Validation simulations are run with default hyperparameters, except for a smaller box size and a lower initial copy number. RMSD is calculated only for the succeeded cases between the COM of design model and the NERDSS assembled structure. (see FigureS3)
